# Increased tissue tension caused by depletion of CLDN3 in the non-neural ectoderm causes neural fold fusion defects in chick embryos

**DOI:** 10.1101/2025.10.14.682463

**Authors:** Elizabeth-Ann Legere, Marie Dumont, Yojiro Yamanaka, Gabriel L. Galea, Aimee K. Ryan

## Abstract

Neural tube morphogenesis provides a dynamic setting in which to study epithelial cell behaviours. Members of the claudin family of tight junction proteins regulate apical epithelial cell behaviors at all steps of neural tube development. We discovered that CLDN3, expressed in the non-neural ectoderm but not the neural ectoderm, is required to mediate neural fold fusion in chick embryos, particularly in the spinal region of the embryo. Depleting CLDN3 affects apical protein localization and apical domain morphology. Here, we used live imaging to re-examine the process of neural fold fusion in the cranial and spinal regions of the embryo and assessed biomechanical parameters of the non-neural ectoderm that are dependent on CLDN3. Our live imaging confirmed previous reports that unlike neural fold fusion in the cranial region and posterior neuropore, the spinal region does not depend on progressive fusion driven by “zippering” cell behaviors but instead fuses in a multi-step process where contact occurs simultaneously at multiple points along the anterior-posterior axis. We and others refer to the process of spinal neural fold fusion as “buttoning” to highlight the differences in cell behaviors from those observed during zippering (van Straaten et al., 1993). CLDN3-depletion decreased the rate of progression of neural fold buttoning within the spinal region. Using cell segmentation analyses we confirmed that CLDN3 depletion decreased the apical cell area of cells at the edges of the neural folds but not of lateral cells in the non-neural ectoderm. CLDN3 depletion increased pMLC staining within the apical domain of the cell, coinciding with a decrease in cell area, suggesting increased epithelial tension. Laser ablation studies revealed that the non-neural ectoderm of CLDN3-depleted embryos exhibits higher tension during neural fold fusion. We showed that treatment with the myosin II inhibitor blebbistatin is sufficient to partially rescue the neural fold fusion defects in CLDN3-depleted embryos. This work provides further evidence for the importance of non-neural ectodermal tissue tension in neural fold fusion and suggests that loss of CLDN3 may alter tissue tension through cytoskeletal regulation pathways within the apical domain. This work supports that CLDN3 contributes to neural fold fusion and epithelial tissue tension during neural fold fusion via modifications to the apical cytoskeleton.

**Summary:** We found that the tight junction protein Claudin-3 (CLDN3) plays a role in regulating tissue tension, by directing actomyosin contraction and apical cell shape/size changes essential for chick neural fold fusion.

## Introduction

Neural tube morphogenesis involves the coordination of complex tissue morphogenetic events that fold a flat epithelial tissue layer into a 3D tube, which is essential for proper central nervous system development. Although the specific cell and tissue behaviors differ between different organisms, the overall steps of neural tube development are conserved (Lee and Nagele, 1985; Sadler, 2005). In chick embryos, neural tube development begins with the formation of the neural plate, a thickening of the neural ectoderm tissue at the anterior midline of the embryo at ∼20hrs of incubation (Hamburger-Hamilton stage (HH) 4) (Hamburger and Hamilton, 1951; Schoenwolf and Alvarez, 1989). The neural plate undergoes convergent extension movements to lengthen along its anterior-posterior axis and narrow along its medial-lateral axis (Schoenwolf and Alvarez, 1989; Wallingford et al., 2002). Tissue at the boundary of the neural ectoderm (NE) and non-neural ectoderm (NNE) elevate dorsally to form the neural folds (NFs) that then converge towards the midline of the embryo, until the opposing neural folds meet and fuse (Lawson et al., 2001; Smith and Schoenwolf, 1989). In chick, the initial site of neural fold contact occurs at the future mid-brain region of the embryo at stage HH8 (∼30hrs) and neural fold fusion continues until stages HH12 or HH14 (∼48-52hrs) in the posterior neural tube (Lawson and England, 1998). Upon fusion, the non-neural ectoderm will separate from the neural ectoderm to form a continuous epithelial layer of non-neural ectoderm over the closed neural tube. This work uses the CLDN3-depleted chick embryo model of neural fold fusion defects to further investigate the process of neural fold fusion, the least understood step in neural tube development (Legere et al., 2024).

Many epithelial cell behaviors in the neural and non-neural ectoderm are required for neural tube development, including cell neighbour exchanges during the intercalation events of convergent extension, changing apical domain size during the formation of hinge points required for neural fold elevation, and in several species, but not chick, formation of membrane protrusions from the non-neural ectoderm that stabilize contact between fusing neural folds (Escuin et al., 2015; Kinoshita et al., 2008; Legere et al., 2024; Wallingford et al., 2002). Each cell in the neural and non-neural ectoderm is connected to its adjacent neighbours by junctional complexes. Tight junctions demarcate the boundary between the apical and lateral membrane domains and are involved in cell-cell adhesion, maintenance of apicobasal polarity, and regulation of paracellular solute flow (Farquhar and Palade, 1963; Matter and Balda, 2003). The evolutionarily-conserved claudin family of tetraspanins is essential for tight junction function through intra- and intercellular interactions with other claudins (Furuse et al., 1998; Krause et al., 2008; Mukendi et al., 2016). Their C-terminal cytoplasmic domains bridge the tight junction to cytoplasmic proteins via scaffolding proteins, including ZO1, and have been shown to functionally interact with the cytoskeleton, apical polarity proteins, adhesion proteins, post-translational modifying proteins, and transcription factors (Hagen, 2017; Suarez-Artiles et al., 2022; Tanaka et al., 2005; Van Itallie et al., 2017). Similarly to adherens junctions, tight junctions can interact with cortical actin networks, a hallmark of the apical domain of epithelial cells (Van Itallie et al., 2017). Regulation of cortical actin is a major mechanism underlying almost all cell morphogenetic behaviors, providing a link between claudins and actin-dependent cell behaviors during neural tube development (Escuin et al., 2015; Van Straaten et al., 2002; Zhou et al., 2020).

The molecular and cellular events required for the early phases of neural tube closure are relatively well understood. However, significantly less is known about mechanisms underlying neural fold fusion, which completes closure of the neural tube at that axial level. Fusion can be broadly defined to encompass the developmental window between elevation of the neural folds and the completion of the cellular remodelling events that give rise to the continuous layer of non-neural ectoderm that overlies the closed neural tube. It can be subdivided into three phases: convergence of the neural folds to the dorsal midline, stabilization at points of contact, and closure/remodelling of cell-cell junctions to unite the two sides of non-neural ectoderm into a continuous layer overlying the closed neural tube. Of the many unanswered questions about neural fold fusion, one area of increased focus is the involvement of the non-neural ectoderm in this process. Initially it was that the non-neural ectoderm creates a pushing force during neural fold convergence (Alvarez and Schoenwolf, 1992), while other data suggest that the non-neural ectoderm moves passively in response to forces generated by the neural ectoderm (Christodoulou and Skourides, 2022; Sokol, 2016), or that it may even act as an apposing force to fusion (Christodoulou and Skourides, 2022; Nikolopoulou et al., 2019).

In mice, apical membrane protrusions arising from the non-neural ectoderm are hypothesized to be involved in maintaining contact between fusing neural folds, however similar protrusions are not seen in chick embryos (Legere et al., 2024; Rolo et al., 2016). After initial neural fold contact, fusion occurs progressively, driven by movement of the closure point where active fusion is occurring. This method of fusion is called “zippering” and was found to be achieved by sequential cell junction shortening in *Ciona* (Hashimoto et al., 2015). In mice, cells at the zippering point form a semi-rosette structure and are dependent on interactions between integrins and the basal extracellular matrix to drive zippering, in combination with constriction of an actomyosin cable at the junction of the non-neural ectoderm and neural ectoderm (Molè et al., 2020; Zhou et al., 2020). The chick embryo does not display an actomyosin cable between the non-neural ectoderm and the neural ectoderm, suggesting that an alternative mechanisms must be driving neural fold fusion in chick embryos (Maniou et al., 2024; Pérez-Verdugo et al., 2025).

In contrast to mouse embryos, which are still U-shaped at the onset of neural fold fusion, most vertebrate embryos, including chick, humans, and rabbit embryos, form three flat germ layers. Thus, it is expected that there must be differences in how local tissue force is generated and transmitted during neural tube development (Peeters et al., 1998a). Indeed, chick embryos also have different patterns of closure points compared to mice, with initial contact between neural folds occurring at the midbrain region of the embryo. Closure then progresses anteriorly until the anterior neuropore (ANP) is closed and posteriorly to an opening between the hindbrain and spinal boundary, also known as the rhombocervical neuropore (RCNP). Between the RCNP and the posterior neuropore (PNP), the neural folds meet at multiple instances, referred as button-like or multistep closure (Pérez-Verdugo et al., 2025; Van Straaten et al., 1996; van Straaten et al., 1993; Wang et al., 2017). Further research investigation of the biomechanics in chick neural fold fusion, along with an understanding of biomechanical regulators is needed.

Our interest lies in the roles of claudin-based tight junctions in neural tube closure. Claudin expression is temporally and spatially regulated during all stages of development, including neural tube morphogenesis (Collins et al., 2013). Manipulating claudin expression in the non-neural or the neural ectoderm provides a mechanism to independently examine the effect of each tissue in chick neural tube development. Previous research from our lab identified a role for CLDN8 and CLDN4 in regulating planar cell polarity proteins in the neural ectoderm during convergent extension and neural fold elevation (Baumholtz et al., 2017). Additionally, the localization of the actin modifying proteins CDC42, RHOA, and pMLC are affected in the neural ectoderm of CLDN4/8-depleted embryos. NTDs are also observed when CLDN3, which is specifically expressed in the non-neural ectoderm, is depleted (Legere et al., 2024). CLDN3-depleted embryos have NTDs due to a failure in neural fold fusion, with the largest proportion of NTDs in the future spinal region of the embryo (Legere et al., 2024). Cells within the non-neural ectoderm of CLDN3-depleted embryos have smaller apical areas and increased number of membrane blebs, a sign of unstable cytoskeletal to membrane attachments. In addition, within the apical domain, there is an increase in the apical polarity proteins PAR3 and MPP5, and a decrease in bicellular localized F-actin compared to control embryos.

As described above, there is a lack of knowledge about the molecular mechanisms underlying biophysical forces regulating chick neural fold fusion. As cell adhesion proteins, including claudins in tight junctions, are linked to the regulation of biomechanical properties of epithelial cell layers, we are examining potential roles of CLDN3 in the regulation of non-neural ectodermal tension during neural tube closure (Chu et al., 2018; Morita et al., 2012). In this study we first revisited the process of chick neural fold fusion using live-imaging and confirmed that the process by which the neural folds converge, and fuse is different from the zippering process observed within the chick cranial region and posterior neuropore. Depleting CLDN3 from the non-neural ectoderm during this process revealed that CLDN3 has a role in regulating the rate of closure and several characteristics of the non-neural ectoderm including cell size, epithelial tension and pMLC expression. We observed that chick spinal neural folds simultaneously converge at multiple regions leaving gaps between the contact points that will fuse shortly after. We and others have termed this process “buttoning”. We also identified increased pMLC signal, suggesting increased actomyosin contraction within the apical domain of CLDN3-depleted non-neural ectodermal cells. CLDN3-depletion also increased tissue tension within the non-neural ectoderm, suggesting that CLDN3 regulates tissue tension during neural fold fusion to mediate proper convergence. This idea is further supported by the fact that the myosin II inhibitor blebbistatin was able to partially rescue neural tube defects arising from CLDN3 –depletion. This work highlights the importance of tissue tension in neural fold fusion and the role that cell junctional proteins, such as CLDN3, play in regulating apical cytoskeletal dynamics.

## Results

### The process of neural fold fusion is distinct between cranial and spinal regions

Previous studies suggest that there are multiple points of closure down the future spinal region of the chick embryo (van Straaten et al., 1993). To study this in real time, we used live imaging and focused on the boundary between the cranial and spinal region. The first point of contact between neural folds occurred at the future midbrain (MB) region at HH8 (∼30h) and then the opposing neural folds progressively closed anteriorly and posteriorly through a zippering action (Fig. 1A, T0-T3). Upon reaching the future hindbrain region, the posterior-directed zippering stalled, and the neural folds bulged at the level of the first somite and contact was made between the opposing neural folds to form the posterior boundary of the rhombocervical neuropore (RCNP) (Fig. 1A, T3-T4). Our *ex vivo* live-imaging data showed that the RCNP forms at ∼HH9 and occurs from the hindbrain with its posterior boundary at the level of somite 1 in most control embryos (n= 3/4) and at the level of somite 3 in 1 embryo. We observed that the length of time it took for the RCNP to close ranged from 30-55 minutes (data not shown). We do not know if the same variability occurs *in ovo*.

**Figure 1.**
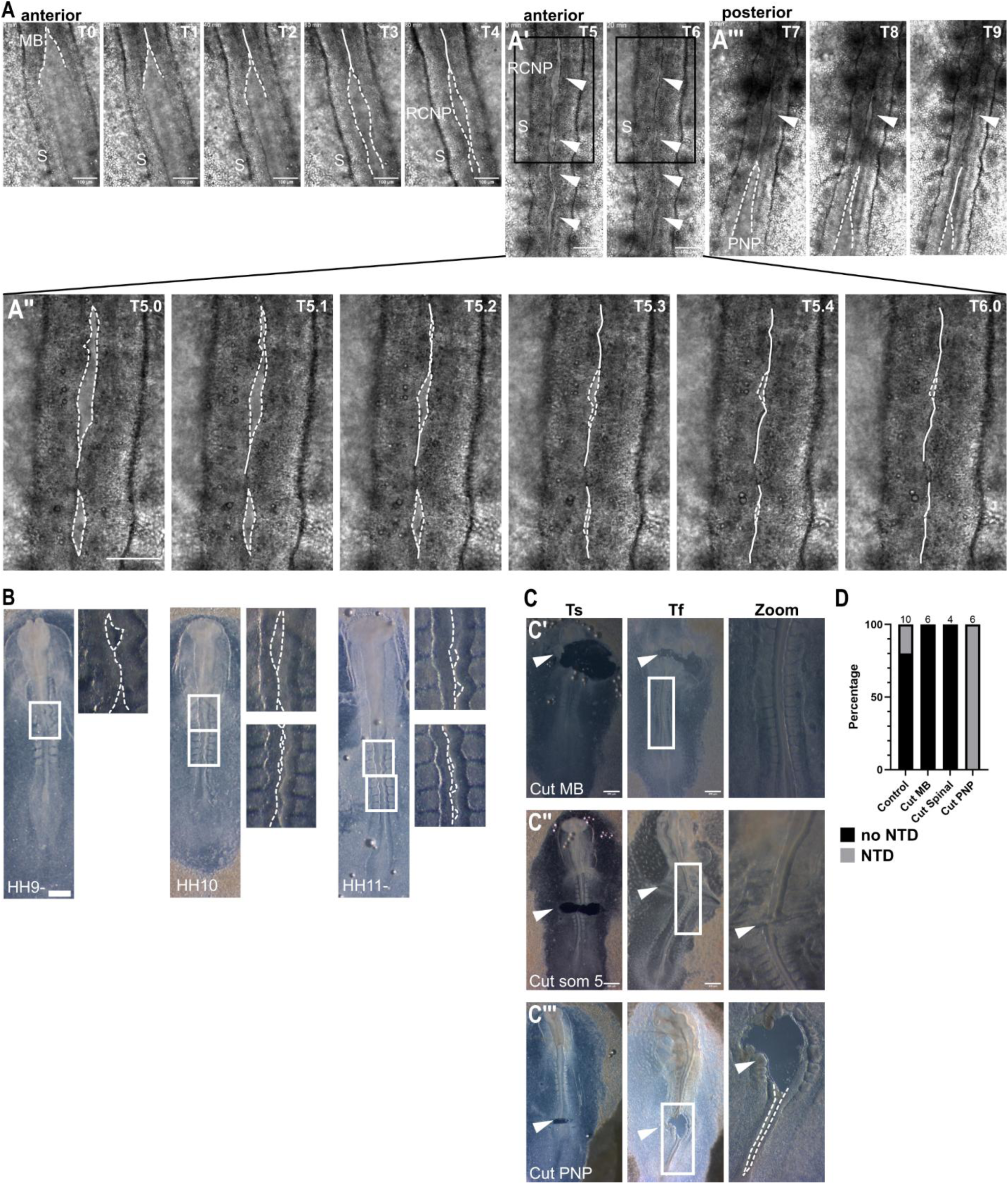
The process of neural fold fusion is distinct between cranial and spinal regions. A) Representative maximum intensity projections (MIPs) from live-imaging of control (GST; n=5 embryos at 20 min intervals (T0-T9)). Midbrain (MB), 1^st^ somite (S), rhombocervical neuropore (RCNP) and posterior neuropore (PNP) are labeled in white. White dotted lines outline the edges of zippering neural folds (NF), and white solid lines outline the edges of fused (by zippering) NFs. White arrowheads mark button gaps. A’) Imaging was paused to adjust focus. Black boxes mark regions shown in higher magnification still in A’’) 5 min intervals (T5.0-T6.0). A’’’) Imaging was stopped to readjust the embryo’s position to image the posterior region of the future spine. B) Representative images of fixed embryos imaged under brightfield at stage HH9 (n=17), HH10 (n=12), and HH11(n=11). White boxes show regions imaged in the corresponding higher magnification images. White dotted lines trace the edges of the NFs. C) Representative images of embryos that were cut at stage HH8 (C’) at the first NF contact point (n=6), at stage HH9 (C’’) at the level of ∼ somite 5 in the mid-spine region (n=4), or at HH11 (C’’’) through the anterior region of the PNP (n=6). Images were taken immediately after cutting (Ts) and after culturing (>15hrs) (Tf). White boxes outline regions shown in the higher magnification images image (Zoom). White arrowheads mark cut, and white dotted lines outline open NFs in PNP. E) Quantification of the proportion of neural tube defects (NTDs) in control (uncut) and cut embryos. Black = no NTD, grey = NTD. Scale bars: A) 100µm, B) 50µm, C) 200µm

Almost simultaneously, multiple additional smaller bulges along the anterior-posterior axis of the spinal NFs folds formed rapidly to create multiple contact points. The gaps between these contacts then appeared to close independently within the next 20-30 minutes (Fig. 1A’, T5-T6,A’’). The most posterior contact point formed the anterior boundary of the posterior neuropore (PNP) and from this point the neural tube fused with stereotypical posterior-directed zippering (Fig. 1A’’’, T7-T9). We confirmed that the RCNP and multiple openings along the spinal region of the embryo were present in both *ex ovo* (Fig. 1B) and *in ovo* cultured embryos (Supp. Fig. 1A), and that both the non-neural ectoderm and the neural ectoderm were unfused within the spinal gaps (Supp. Fig. 1B). This process of spinal specific neural fold fusion has previously been referred to as “multistep,” or “button-like” (Lawson and England, 1998; van Straaten et al., 1993). We observed that during spinal neural fold fusion the neural folds have the appearance of a shirt being stretched to create gaps in between the buttons. Going forward we will refer to the spinal neural fold fusion as buttoning, with the openings between buttons called gaps. The RCNP is the boundary between the cranial zippering and buttoning observed in the spinal region.

By combining data obtained from live imaging and fluorescence confocal images of ZO1 signal to view cell outlines, we defined three steps that occur during buttoning: convergence, contact, and fusion. We observed that beginning at HH8 the elevated unfused neural folds appeared “wavy” and periodically bulged out (Fig. 1A, B; HH9-). Our data from phalloidin-stained embryos agreed with previous observations that chick embryos do not have a supracellular actin cable between the non-neural and the neural ectoderm and also suggested that bulging of the neural folds is not driven by specific actomyosin constriction at the neural edge, as there is no enrichment of F-actin at unfused neural fold edges (Maniou et al., 2024) (Supp. Fig. 1D). Following the formation of the RCNP (∼HH9) the spinal neural folds button. The gaps between buttons were noticeably smaller than the RCNP and usually occurred between somites (Fig. 1A’, B (HH10), Supp. Fig. 1C). We observed that these gaps appeared to close progressively from both the anterior and posterior direction until a small area in the center remained (Fig. 1A’’). The whole-mount immunofluorescence images of ZO1 signal suggested that the neural folds came together in the gaps (Supp. Fig. 1C1), and that the last opening was fused when the cells pinched together (Supp. Fig. 1C2) to form a rosette-like shape (Supp. Fig. 1C3, D’).

To determine if zippering and buttoning involved distinct cell behaviors, we performed experiments previously used to evaluate the zippering of neural folds in *Ciona intestinalis* (*Ciona*). During *Ciona* neural tube development, the neural folds contact each other and then progressively zipper together in a process that is driven by junctional shortening at the zippering point (Hashimoto and Munro, 2019). When researchers cut *Ciona* embryos in half during the process of neural fold zippering, they observed that the half that contains the point of contact (zippering point) would complete neural fold fusion as normal, however the half that did not contain the zippering point would not (Hashimoto et al., 2015). To determine if chick embryos depend on progressive junctional shortening to drive zippering as described in the *Ciona* or to form the semi-rosette shape at the zippering point as described in the mouse (Molè et al., 2020), we adapted this method for the chick embryo to test whether the zippering and buttoning processes use different cell behaviors. Briefly, we cultured chick embryos until the appropriate stage, removed the vitelline membrane, and then used a hair-knife to cut the embryos at one of three locations: posterior to the first closure point in the midbrain, at the level of somite 5, or in the posterior neuropore. Cut embryos were cultured dorsal side-up for up to 24hrs and assessed for neural tube closure (Fig 1C). Embryos cut posterior to the first closure point in the midbrain at HH8 were assessed to determine if the RCNP would form and if the spinal neural folds would fuse. The spinal regions in all these embryos were able to fuse and complete neural tube closure (n=6) (Fig. 1C’). In a second series, we assessed the effect of cutting through the forming neural tube at the level of the 5^th^ somite in HH9 embryos to assess the effect on the process of spinal neural fold fusion. Again, the spinal regions of all these embryos were able to complete neural fold fusion (n=4) (Fig. 1C’’). These data suggest that unlike the zippering in *Ciona* that depends on the behavior of cells at the zippering point, the chick neural folds can fuse at the button points independent of the presence of the zippering point or continuous zippering. To confirm that zippering would not progress without the presence of a zippering point, we cut older (HH10/11) embryos at the anterior region of the posterior neuropore (PNP) and allowed them to develop until they reached HH14 to determine if fusion of the PNP occurred. All cut embryos contained neural fold fusion defects in the PNP (n=6) compared to 50% of uncut control embryos that had their vitelline membrane removed (n=2/4) (Fig. 1C’’’).

### CLDN3-depletion decreased the rate of neural fold convergence during fusion

Given that the NTDs in CLDN3-depleted embryos occur primarily in the future spinal region (Legere et al., 2024), we examined whether there were visible differences during spinal neural fold fusion between control and CLDN3-depleted embryos. Previously we found that CLDN3-depleted embryos form neural fold hinge points and do not have a delay in closure of the PNP, but we did not examine which step of neural fold fusion was affected (Legere et al., 2024). To be able to image embryos midway through neural fold fusion (HH8-HH10), embryos were cultured *ex ovo* on agar-albumen plates (Chapman et al., 2001) containing either the GST-C-CPE variant GST-C-CPE^LDR^ (LDR) µg/mL) that binds and removes CLDN3 from tight junctions or molarity matched GST (266 µg/mL) (Legere et al., 2024; Veshnyakova et al., 2012). We first measured the area and perimeter of RCNPs of stage matched HH9-/HH9 control (GST) (n=4) and CLDN3-depleted (LDR) (n=5) embryos and found that on average CLDN3-depleted embryos had larger RCNP areas (GST: 448.0 µm^2^ ± 354.5, LDR: 2802.6 µm^2^ ± 979.1, p = 0.003) and perimeters (GST: 117.2 µm ± 22.0, LDR: 277.0 µm ± 46.5, p = 0.0005). These results confirmed that the effect of CLDN3 depletion on spinal neural fold fusion was occurring during the process of neural fold convergence in the progression of buttoning.

To visualize the formation of the RCNP in the CLDN3-depleted embryos we performed live imaging on chick embryos cultured on either LDR (400 µg/mL) or molarity matched control GST (266 µg/mL) agar-albumin plates. In a second series, all control embryos (n=5/5) completed neural tube closure in the 15 hours that the embryos were observed with live imaging. In contrast, only half of the CLDN3-depleted embryos (n=2/4) had closed neural tubes. There was no effect on the rate of zippering in the midbrain region of the CLDN3-depleted embryos relative to the control embryos (GST: 2.7 µm/min ± 1.4, LDR: 2.5 µm/min ± 1.0, p = 0.79) (Fig. 1A, Fig. 2D). To compare the rate of RCNP neural fold closure between control and CLDN3-depleted embryos, we determined the change in the distance between neural folds at the midpoint of the RCNP over time. On average the rate of RCNP closure was slower in CLDN3-depleted embryos (GST: 1.9 µm/min ± 0.7, LDR: 0.6 µm/min ± 0.3, p = 0.01) (Fig. 2D). These results suggest that the larger RCNP in CLDN3-depleted embryos was due to an overall slowing of the rate of neural fold convergence, a process that occurs throughout buttoning and may explain why CLDN3-depleted embryos have spinal NTDs (Legere et al., 2024).

**Figure 2.**
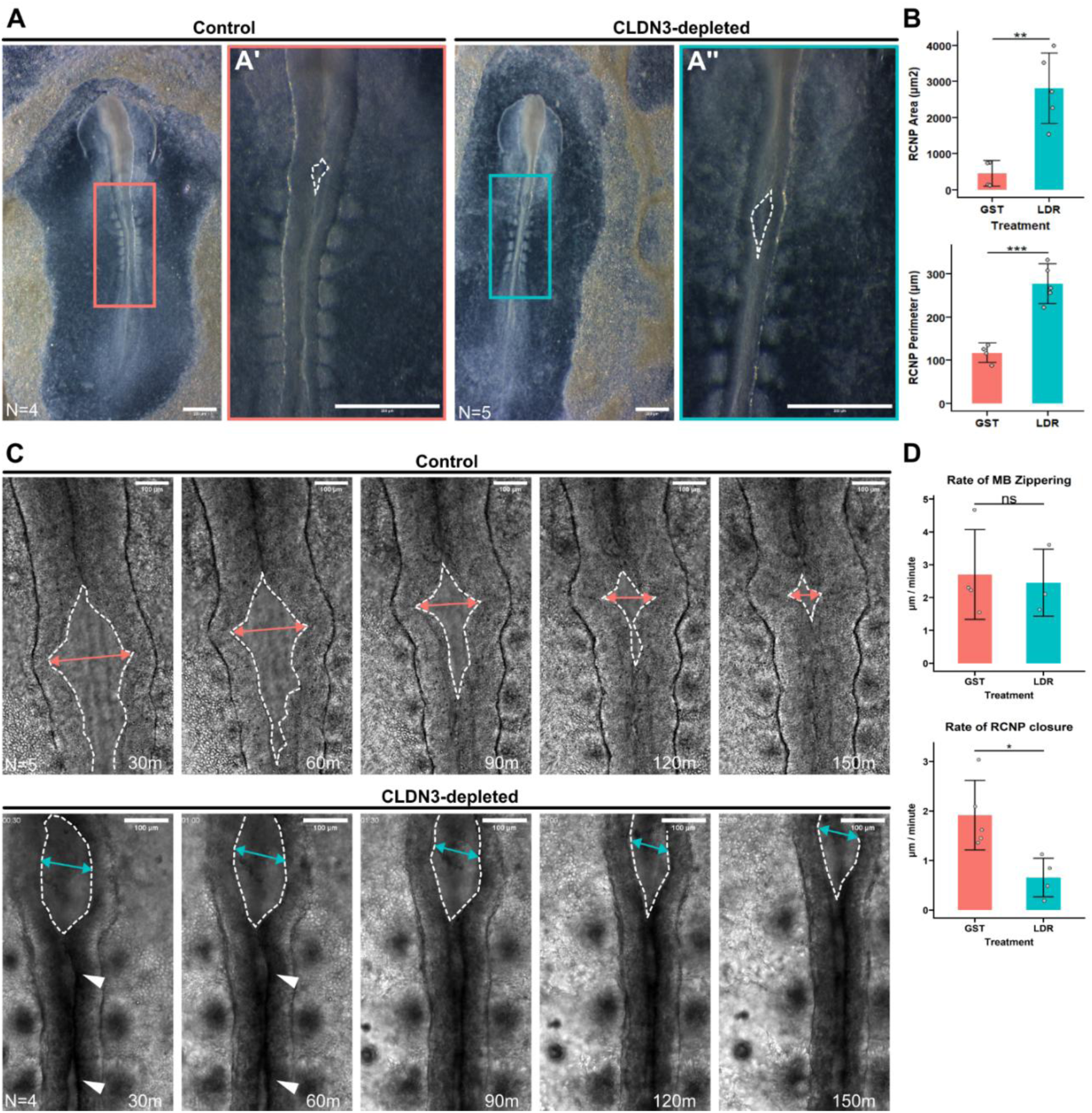
CLDN3-depletion decreased the rate of NF convergence during fusion. A) Representative images of control (GST; n=4) and CLDN3-depleted (LDR; n=5) embryos at stages HH9- and HH9. Coloured boxes illustrate region of image shown at higher magnification on right; white dotted lines outline the rhombocervical neuropore (RCNP). B) Quantification of RCNP area and perimeter. Coloured bars (GST: pink, LDR: blue) show the mean; each point is the measurement from one embryo. Error bars show standard deviation. C) Maximum intensity projections (MIPs) of still images from live-imaging of control (n=5) and CLDN3-depleted (n=4) embryos taken at 30-minute (30m) intervals. White dotted lines outline the RCNP and coloured lines (GST: pink, LDR: blue) mark measured distance between RCNP. White arrowheads point out regions that are open between NFs. D) Quantification of the rate of change of zippering in the midbrain (MB) and quantification of rate of change between fusing RCNP NFs. Coloured bars (GST: pink, LDR: blue) show the mean, and each point is the measurement of one embryo. Error bars show standard deviation. Statistical significance was evaluated with Student’s T-test. *** p < 0.005, ** p < 0.01, * p< 0.5. Scale bars: A) 200µm, C) 100µm.

### Apical cell morphology is heterogeneous in the non-neural ectoderm

In our previous analysis of the effects of CLDN3 depletion on apical cell morphology, we manually traced the outline of a limited number of cells from scanning electron micrographs and found that on average the apical cell areas of the CLDN3-depleted embryo were smaller (Legere et al., 2024). To systematically measure cell parameters on a larger cell population, we segmented cells in the non-neural ectoderm using images of ZO1 fluorescence at the level of the first somite (Fig. 3A,B) and compared apical cell measurements between control (n=4 embryos, N_­_= 2948) and CLDN3-depleted (n=4, N_­_= 2974) embryos. There was no significant difference in the average cell area between control (52 µm^2^ ± 36) and CLDN3-depleted (48 µm^2^ ± 30) embryos (Supp. Fig. 2A).

**Figure 3.**
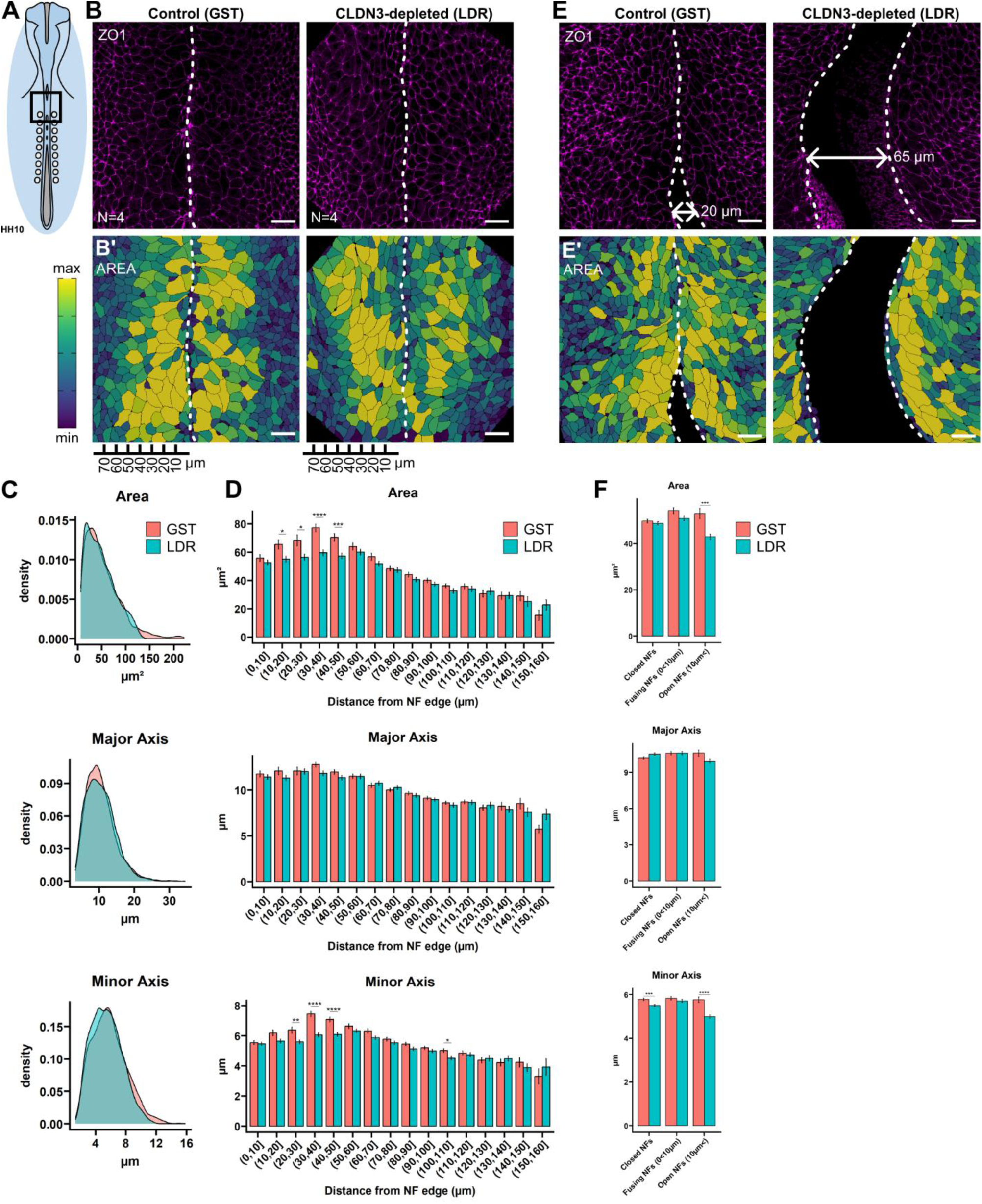
Apical cell morphology was heterogeneous in the non-neural ectoderm. A) Schematic of an HH10 chick embryo. The black box outlines the approximate region imaged. B) Representative surface peeled images of control (n=4) and CLDN3-depleted (n=4) embryos after immunofluorescence with ZO1 antibody (magenta). White dotted lines outline the edges of the neural folds (NFs). B’) Images were segmented using ZO1 signal and apical surfaces were coloured based on apical cell area (yellow = largest, purple = smallest). White dotted lines outline the edges of the neural folds (NFs). The ruler below the images illustrates the distance from the NF edge in 10µm increments. C) Density plots of cell measurements from segmentation analysis of area (µm^2^), major axis (µm), and minor axis (µm) of GST control (pink) (N_­_= 2948) and LDR CLDN3-depleted (blue) (N_­_= 2974). D) Analysis of cell area (µm^2^), major axis (µm), and minor axis (µm) of cells binned by distance of the cell centroid from the edge of the NF. Bars show the mean; error bars show standard error of GST control (pink) and LDR CLDN3-depleted (blue). E) Representative surface peeled images of GST control and LDR CLDN3-depleted embryos showing an example for each where the NFs are open. White dotted lines outline the edges of the neural folds (NFs). An example distance between open NFs is shown with the bidirectional arrow. E’) Cell areas from segmented images were coloured (yellow = largest, purple = smallest). White dotted lines outline the edges of the neural folds (NFs). F) Quantification of cell area (µm^2^), major axis (µm), and minor axis (µm) binned by the distance between NFs. Closed NFs = 0µm (GST: N_­_= 1540, LDR: N_­_= 1607), Fusing NFs = 0µm <= 10µm (GST: N_­_= 1083, LDR: N_­_= 780), and Open NFs = > 10µm (GST: N_­_= 320, LDR: N_­_= 587). Bars show the mean; error bars show standard error of GST control (pink) and LDR CLDN3-depleted (blue). D, F) Statistical significance between GST and LDR in each bin was assessed with Student’s T-test with multiple comparisons p-value correction with Benjamini-Hochberg (False Discovery Rate). **** p < 0.001, *** p < 0.005, ** p < 0.01, * p< 0.5. Scale bars: 20µm

When the segmented cells were coloured according to their area, a pattern emerged suggesting that cells closer to the neural folds were larger than cells in more lateral non-neural ectoderm in control embryos (Fig. 3B’). To quantify this based on the location of cells within the non-neural ectoderm, we binned cells based on their distance from the neural fold edge in 10 µm intervals and analyzed their apical cell areas. There were comparable numbers of control and CLDN3-depleted cells within each bin. In control embryos, we found that the cells closest to the edge of the neural fold (0 - 10µm) were smaller than the cells from bins between 20µm and 50µm (GST 0-10µm: N_­_=266, 56 µm^2^ ± 38; GST 20-30µm: N_­_=158, 68 µm^2^ ± 48; GST 30-40µm: N_­_=156, 77 µm^2^ ± 34; GST 40-50µm: N_­_=192, 70 µm^2^ ± 36), but larger than cells in bins that were between 80 and 140µm from the edge of the neural fold (GST 80-90µm: N_­_=312, 44 µm^2^ ± 29). In contrast, CLDN3-depleted cells at the edge of the neural fold (0 - 10 µm) were not smaller than cells in bins of 20–50µm, but were larger than cells in bins between 80 and 150µm (LDR 0-10µm: N_­_=281, 53 µm^2^ ± 35; GST 30-40 µm: N_­_=227, 60 µm^2^ ± 32; GST 80-90µm: N_­_=259, 41 µm^2^ ± 25) (Supp. Fig 2B). We found that the apical areas of cells from CLDN3-depleted embryos with a centroid 10-50µm away from the neural fold edge were significantly smaller than cells the same distance from the neural fold edge in control embryos. The maximum difference between the apical surfaces in control and CLDN3-depleted embryos was for cells with centroids 30-40µm away from the neural fold edge (GST: N_­_= 156; 77 µm^2^ ± 34, LDR: N_­_= 227; 60 µm^2^ ± 32, p = 0.000008) (Fig. 3D).

To examine whether this effect was related to the stage of neural fold closure, we binned the cells intro three categories: closed neural folds (0µm between the left and right neural fold edges), fusing neural folds (0 ≤10µm), and open neural folds (>10µm) (Fig. 3E, E’). In control embryos, cells at closed neural folds were smaller than cells at fusing neural folds (NFs) (GST closed NFs: N_­_= 1540; 50µm^2^ ± 31; GST fusing NFs: N_­_=1083; 54µm^2^ ± 42, p=0.005) while in CLDN3-depleted embryos, cells at open neural folds were smaller than cells at fusing and closed neural folds (LDR open NFs: N_­_=587, 43µm^2^ ± 29; LDR fusing NFs: N_­_=780, 51µm^2^ ± 30; LDR closed NFs: N_­_=1607, 49µm^2^ ± 30, p=0.000003, 0.0002) (Supp. Fig. 2B’). Only cells from LDR-treated embryos that had centroids in line with open neural folds (> 10µm) were significantly smaller compared to control cells in the same bin (GST: N_­_=325, 53µm^2^ ± 40; LDR: N_­_=587, 43µm^2^ ± 29; p=0.0001) (Fig. 3F). Finally, we compared control and CLDN3-depleted cells binned by distance from the neural fold edge and distance between neural folds and found that the smaller CLDN3-depleted cells at open neural folds were largely from cells at the edge of the neural folds (0-10 µm) (GST: N_­_=51, 63µm^2^ ± 39; LDR: N_­_=81, 34µm^2^ ± 27; p=0.0001). CLDN3-depleted cells from closed neural folds were smaller at distances between 10 and 50µm, with a maximum difference at cells with centroids 30-40µm away from the neural fold edge (GST: N_­_=67, 85µm^2^ ± 36; LDR: N_­_=134, 58µm^2^ ± 30; p=0.00001) (Supp. Fig. 2C). In conclusion, CLDN3 depletion led to decreased apical cell areas of cells that were closest to the edge of the neural folds, especially in cells that were bordering open neural folds.

To assess whether the decrease in apical cell area in CLDN3-depleted embryos was due to an overall loss of area, or due to cells changing shape we compared major and minor axis from fit ellipses. There was no change in the major cell axis at any distance from the neural fold between control and CLDN3-depleted embryos (Fig. 3D). However, there was a significant decrease in the minor cell axis in CLDN3-depleted embryos with a centroid 20-50µm away from the neural fold edge, with the largest effect observed from cells with centroids 30-40µm away from the neural fold edge (GST: N_­_=156, 7.4µm ± 2.2; LDR: N_­_=227, 6.0µm ± 2.1; p = 0.00000002) (Fig. 3D). Minor axes of cells from CLDN3-depleted embryos were smaller independently of whether the neural folds were fused (0µm) (GST: N_­_= 1540, 5.8µm ± 2.0; LDR: N_­_=1607, 5.5µm ± 1.9; p=0.0001437) or open (>10µm) (GST: N_­_=325, 5.8µm ± 2.2; LDR: N_­_=587, 5.0µm ± 1.9; p=0.0000006) (Fig. 3F). There was no difference in major axes of cells from control and CLDN3-depleted embryos when binned by distance between NFs (Fig. 3F). These data suggest that the decrease in apical cell size of cells close to the neural fold edge in CLDN3-depleted embryos was driven by a decrease along the minor axis of cells. The changes in apical cell morphology may underlie the decrease in neural fold convergence, and formation of NTDs, as the largest difference in cell size and shape was observed at cells aligned with open neural folds.

### CLDN3-depletion increases the level of pMLC apical enrichment

To examine downstream effects of CLDN3-depletion that may underlie changes in apical cell morphology, we examined the expression of phosphorylated myosin light chain (pMLC) in the non-neural ectoderm of control (n=9) and CLDN3-depleted (n=13) embryos undergoing neural fold fusion (Fig. 4A). Unlike what is observed in the neural ectoderm (Nishimura et al., 2012), pMLC does not form mediolateral cables in the non-neural ectoderm but instead it is weakly localized throughout the cytoplasmic face of the membrane and the cytoplasm. On average the overall intensity of pMLC compared to the unchanging ZO1 expression (Supp. Fig. 3A) was increased in CLDN3-depleted embryos (GST: 1 ± 1.2, LDR: 2.2 ± 1.7, p = 0.04) (Fig. 4B, C) in both the membrane (GST: 0.8 ± 0.9, LDR: 1.7 ± 1.2, p = 0.04) and cytoplasmic (GST: 1.3 ± 1.6, LDR: 2.9 ± 2.2, p = 0.04) regions of the cells (Fig. 4 C’, Supp. Fig. 3B). As expected due to CLDN3 only being expressed in the non-neural ectoderm, no effect on pMLC intensity was observed in the neural ectoderm of CLDN3-depleted embryos (GST: 2.2 ± 0.9, n=3, LDR: 3.5 ± 2.1, n=6, p = 0.25) (Supp. Fig. 3C). The increased level of pMLC in CLDN3-depleted embryos may partially explain the decreased apical cell areas, since increased levels of pMLC is known to correlate with increased cortical actomyosin constriction and decreased apical domain area (Kinoshita et al., 2008). In CLDN3-depleted embryos, we observed several instances of increased pMLC level on some cells facing the open neural folds (Fig. 4B), suggesting that CLDN3 depletion prevented the relaxation of tension in the closest zone to the neural fold edge.

**Figure 4.**
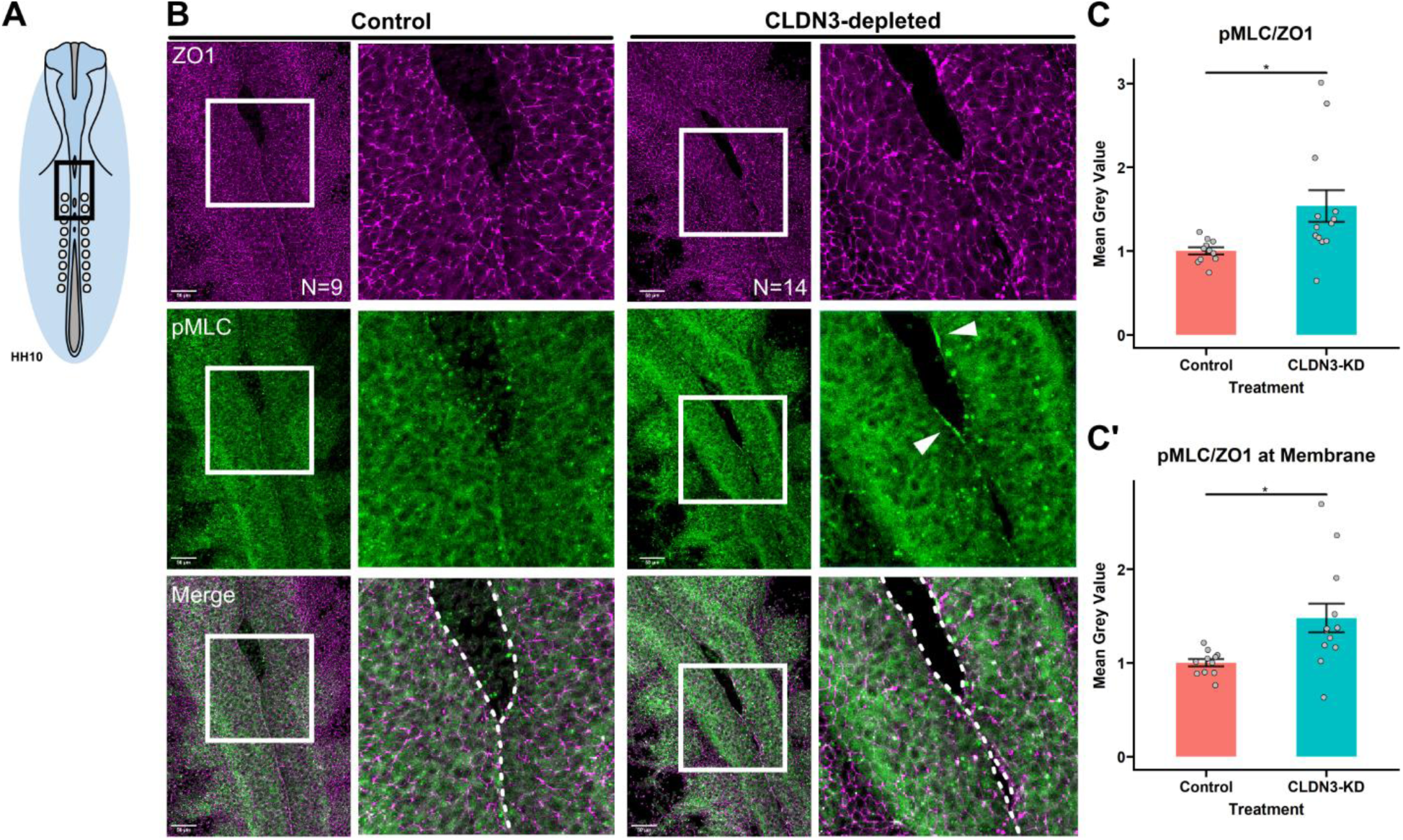
CLDN3-depletion increased expression of pMLC. A) Schematic of a HH10 embryo with the black box showing the approximate area imaged. B) Representative immunofluorescence surface-projected images of control (GST; n=9) and CLDN3-depleted (LDR; n=13) embryos cultured for 15 hrs stained with anti-ZO1 antibody (magenta) and pMLC antibody (green). White boxes illustrate the region of image shown in adjacent zoomed image. White dotted lines outline the edges of the neural folds (NFs) on the merged images. White arrows point to areas of increased pMLC staining on the NF edge. Quantification of pMLC/ZO1 signal intensity normalized per experiment C) over the entire image and C’) at membrane regions defined by ZO1 signal. C, C’) Coloured bars (control: pink, CLDN3-depleted (CLDN3-KD): blue) show the mean, and each point is the measurement of one embryo. Error bars show standard error. Statistical significance was evaluated with Student’s T-test. * p< 0.5. Scale bars: A) 50µm, C, D) 20µm

### CLDN3-depletion increases tension in the non-neural ectoderm during neural tube development

Previous studies in mouse (De La O et al., 2025; Marshall et al., 2022), *Xenopus* (Morita et al., 2012) and chick (Maniou et al., 2024) used laser ablation to infer tissue tension perpendicular to the direction of the ablations in the non-neural ectoderm during neural tube development. To determine if CLDN3-depletion affected the tension in the non-neural ectoderm, we cultured chick embryos on LDR (400 µg/mL) or molarity matched GST (266 µg/mL) agar-albumin plates until HH8. Embryos were stained with CellMask prior to imaging and ablated along a 50 µm long line. To infer tension of the epithelial layer, we measured the recoil, or the change in distance between two points on either side of the ablation before and after ablation. Higher recoil correlates with increased tension, while lower recoil suggests lower tension (Liang et al., 2016). First, we assessed the amount of tension in the neural ectoderm and the non-neural ectoderm. Tension in the neural ectoderm is anisotropic, with mediolaterally (ML) directed cuts having higher recoil than anterior-posteriorly (AP) directed cuts in control embryos (n=9, ML: 4.0µm ± 3.7, AP: 0.8µm ± 0.7, p = 0.02). There was no significant difference in recoil in either direction between control or CLDN3-depleted embryos in the neural ectoderm (GST ML: n=9; 4.0µm ± 3.7, LDR ML: N=5; 4.5µm ± 5.9, p = 0.84, GST AP: n=10; 0.8µm ± 0.7, LDR AP: n=5; 1.2µm ± 1.3, p = 0.47) (Supp. Fig. 4A). In contrast, tension was not anisotropic in the non-neural ectoderm of control embryos (n=10, ML: 3.7µm ± 3.7, AP: 4.1µm ± 3.2, p = 0.65) nor in the non-neural ectoderm of CLDN3-depleted embryos (n=8, ML: 8.4µm ± 4.7, AP: 8.4µm ± 3.8; p = 0.98). However, there was a significant difference in tension in the non-neural ectoderm between control and CLDN3-depleted embryos in both directions, with higher tension being observed in CLDN3-depleted embryos (GST ML: n=10, 3.7µm ± 3.7; LDR ML: n=8; 8.4µm ± 4.7; p = 0.03, GST AP: n=10; 4.1µm ± 3.2, LDR AP: n=8; 8.4µm ± 3.8, p = 0.02) (Fig. 5B). Because non-neural ectoderm tension was not anisotropic, we also compared the combined recoil measurements for both directions and confirmed that the higher recoil observed after ablation was significantly higher in CLDN3-depleted embryos than in control embryos (GST: n=10, 3.8µm ± 2.9; LDR: n=8; 8.7µm ± 2.4; p = 0.001) (Fig. 5C, D).

**Figure 5.**
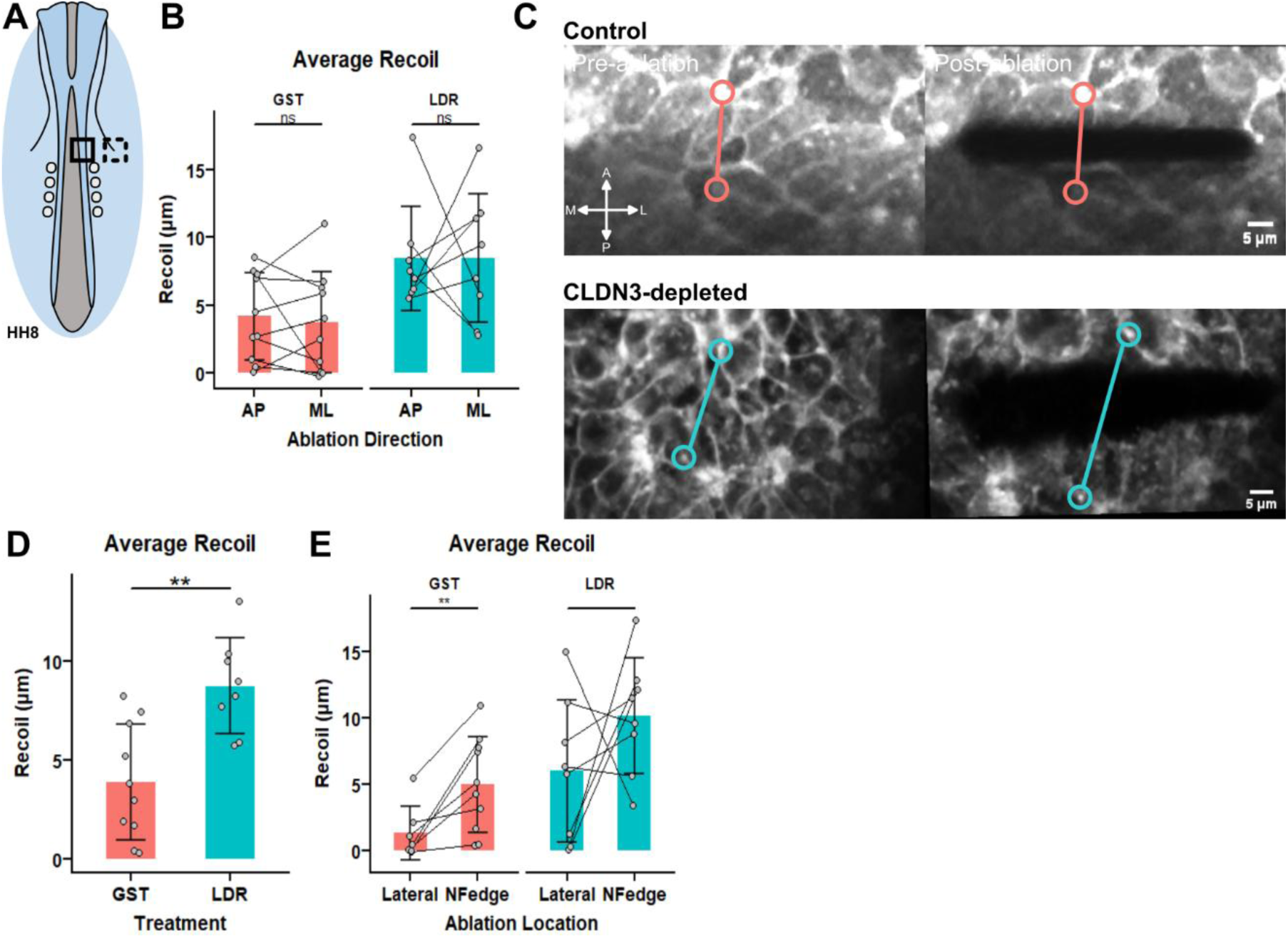
CLDN3-depletion increased tension in the NNE during neural tube development. NNE tension was assessed with tissue-wide (∼5-10 cell wide) laser ablations. Recoil perpendicular to the direction of the ablation was measured by taking the difference in lengths between landmarks pre- and post-ablation. All graphs in B), D), and E) show mean in bars (GST – pink; LDR treated – blue), errors bars show standard deviation, points represent average recoil (µm) per embryo, and lines connect measurements made on the same embryo. A) Schematic of HH8 embryo to illustrate approximate regions ablated on the NF edge (black box) and lateral NNE tissue (black box - dashed). B) Anterior-Posterior (AP) and Medial-Lateral (ML) ablation directions were performed on the same embryo, but on opposing neural folds. AP or ML recoils were averaged per embryo, and paired Student’s T-tests were used to assess difference in recoils done in AP and ML directions in within GST control (n=10; pink), and LDR CLDN3-depleted (n=8; blue). C) Representative images of images used for measurement of recoil of control (n=10) and CLDN3-depleted (n=8), orientation is shown in the lower left corner, and images are labeled “Pre-ablation” and “Post-ablation” from left to right. D) Average recoil of both AP and ML ablations compared between control (n=10; pink) and CLDN3-depleted (n=8; blue). Student’s T-test was used to compare mean recoils. E) Average recoil of AP and ML ablations done within lateral NNE tissue (20 cells from neural fold (NF) edge) and ablations done within cells at the NF edge. Averages from the same embryo are connected by lines and a paired Student’s T-test was used to compare recoil within control (n=7; pink), and CLDN3-depleted (n=8; blue). ns: no significance (p>0.05), ** p < 0.01. Scale bars: 5µm.

To understand whether the increased tissue tension in CLDN3-depleted embryos is limited to specific regions or across the entire non-neural ectoderm, we compared recoil measured following ablations at the neural fold edge with recoil in more lateral regions (∼20 cells away from the edge) (Fig. 5A). In control embryos, the cells at the neural fold edge were under higher tension (n=7, NF edge: 4.9µm ± 3.6, Lateral: 1.3 µm ± 1.9, p = 0.006) while in CLDN3-depleted embryos, the cells at the neural fold edge were not significantly more tense than lateral cells (n=8, NF edge: 10.1µm ± 4.3, Lateral: 6.0µm ± 5.3, p = 0.24). CLDN3-depleted embryos were more tense at both the neural fold edge and in more lateral non-neural ectoderm tissue than in control embryos (NF edge: p = 0.01, Lateral: p = 0.04) (Fig. 5E). Thus, the increase in non-neural ectoderm tension in CLDN3-depleted embryos appears to be related to reduced levels of CLDN3 in tight junctions independently of their proximity to the midline. While the increased tension at the neural fold edge correlates with the smaller apical surfaces observed in CLDN3-depleted embryos, this correlation does not hold in more lateral regions of the non-neural ectoderm.

### Treatment with the myosin II inhibitor blebbistatin partially rescues NTDs in CLDN3-depleted embryos

Given the increase in pMLC and the increase in tension observed in the non-neural ectoderm of CLDN3-depleted embryos, we wanted to determine if reducing the tension would reduce the incidence of NTDs. To do this we used the myosin II inhibitor, blebbistatin to reduce the tension in the non-neural ectoderm in CLDN3-depleted embryos. Blebbistatin treatment was previously shown to reduce tension in the surface ectoderm of mouse embryos at a concentration which does not directly impair progression of neural tube closure (Nikolopoulou et al., 2019). Embryos were cultured on GST control or LDR containing agar-albumen plates and treated with 2.5µM - 50µM blebbistatin at the beginning of neural fold fusion (∼HH8). We observed a dose-dependent effect of blebbistatin on neural tube development in control embryos (Fig. 6 B, B’, Supp. Fig 5A). At the lowest dose tested, 2.5µM blebbistatin, one-third of the control embryos displayed a mild cranial NTD (n=1/3), at 5µM blebbistatin 100% of embryos had NTDs (n=8/8) that varied in severity and location, and at 50µM blebbistatin 100% of embryos displayed NTDs in both cranial and spinal regions (n=3/3) and two-thirds of the embryos (n=2/3) were noticeably shorter along their anterior-posterior axes than untreated embryos (Fig. 6A, Supp. Fig5A). The increased incidence of NTDs in embryos cultured in 5µM blebbistatin-treated versus control media (25% NTDs, n=12) was significant (Fisher’s Exact Test, p = 0.0014). The 5µM blebbistatin dose caused NTDs most similar in penetrance and severity to the NTDs caused by CLDN3 depletion (94% NTDs, n=17, Fisher’s Exact Test to GST, p = 0.0002). When combined with CLDN3 depletion, the low and high doses of blebbistatin had no effect on the incidence of NTDs as compared to CLDN3-depletion alone (400 µg/mL LDR = 94% NTDs, n=16/17; LDR + 2.5µM blebbistatin = 100% NTDs, n=3/3; LDR +50µM blebbistatin = 92% NTDs, n=11/12). However, there is a significant decrease in the number of embryos with NTDs when 5µM blebbistatin was combined with LDR treatment to deplete CLDN3 (LDR + 5µM blebbistatin = 36% NTDs, n=5/14, p=0.0012).

**Figure 6.**
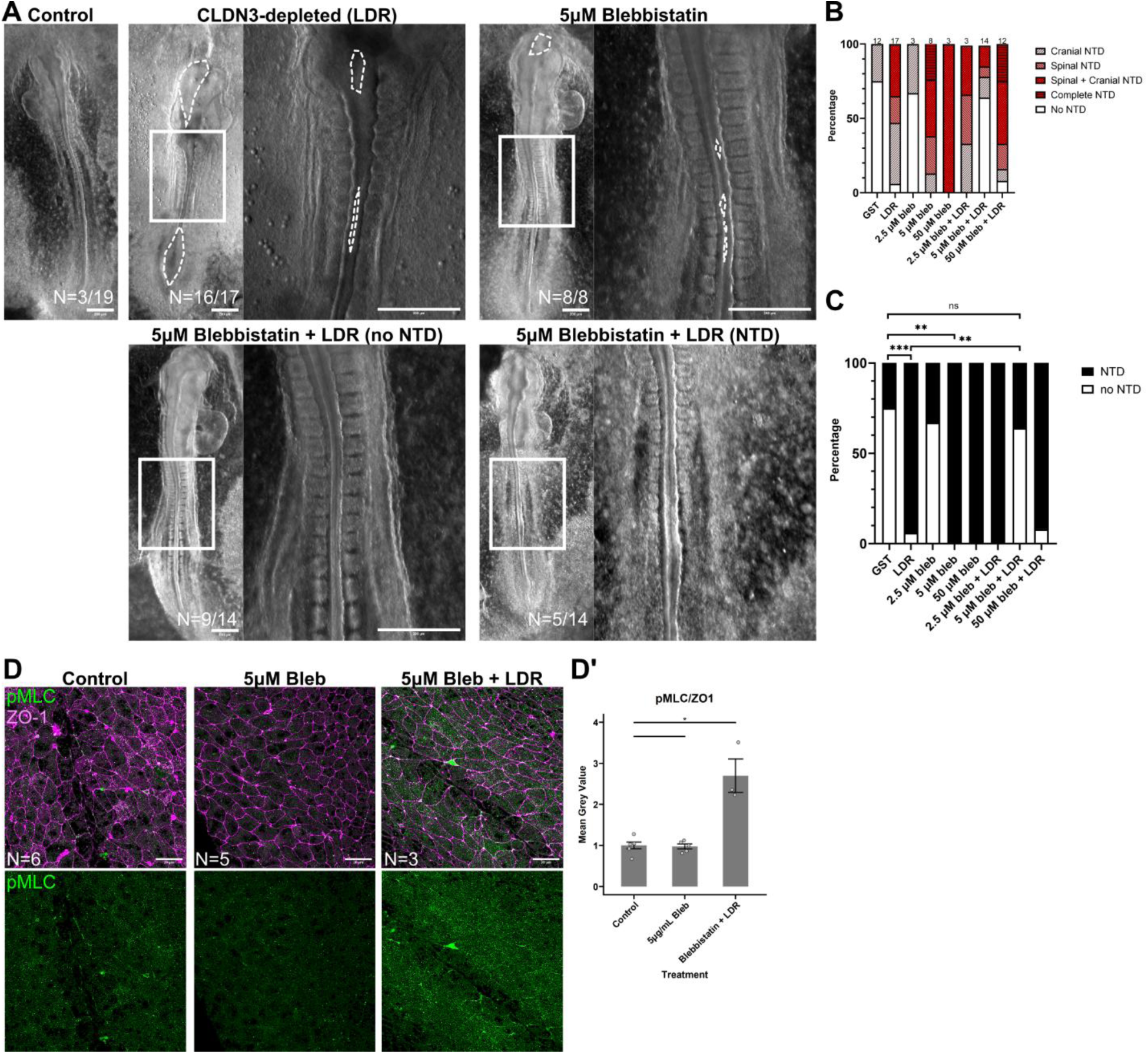
Treatment with myosin II inhibitor blebbistatin rescued the NTDs in CLDN3-depleted embryos. A) Representative images of GST + DMSO control (n=19), 400 µg/mL LDR + DMSO (n=17), 5µM blebbistatin (N=8), 5µM blebbistatin + 400µg/mL LDR (n=14) without a neural tube defect (NTD) (n=9/14) and with NTD (n=5/14) after 24h of culture. White boxes illustrate region of image shown at higher magnification; white dotted lines outline open neural tube defects (NTDs). B) Quantification of NTD location and C) total number of embryos with NTDs. Total number of embryos shown above bars. D) Representative immunofluorescence surface projected images of GST + DMSO control (n=6), 5 µM blebbistatin (Bleb; n=5), and 5µM blebbistatin + 400µg/mL LDR (Bleb + LDR; n=3) cultured for 8h with LDR and 2.5h with Blebbistatin stained with ZO1 (magenta) and pMLC (green). D’) Quantification of pMLC/ZO1 signal intensity in the non-neural ectoderm normalized per experiment. Bars show the mean, and each point is the measurement of one embryo. Error bars show standard error. Statistical significance was evaluated with Student’s T-test. * p< 0.5. Scale bars: A) 200µm, C) 20µm

To determine if the ability of 5µM blebbistatin to rescue NTDs caused by CLDN3 depletion was upstream or downstream of the increased levels of pMLC observed in CLDN3-depleted embryos (Fig. 4), we assessed pMLC expression in control embryos, control embryos treated with 5µM blebbistatin, and CLDN3-depleted embryos treated with 5µM blebbistatin (Fig. 6D). The pMLC/ZO1 ratio in the non-neural ectoderm of 5µM blebbistatin-treated embryos (n=5) was unchanged compared to control embryos (n=6) (GST+DMSO: 0.47 ± 0.09, 5µM blebbistatin: 0.45 ± 0.08, p > 0.05). Similarly to CLDN3-depleted embryos there was a significant increase in pMLC immunofluorescence intensity in the non-neural ectoderm of 5µM blebbistatin + CLDN3-depleted embryos (n=3) compared to embryos treated with only 5µM blebbistatin (LDR + 5µM blebbistatin: 1.32 ± 0.34, 5µM blebbistatin: 0.45 ± 0.08, p = 0.04). Therefore, the partial rescue of CLDN3-depleted NTDs by blebbistatin does not appear to be acting by reducing the levels of pMLC and is most likely acting downstream of this effect or via a parallel pathway due to release of excess non-neural ectodermal tension through inhibition of myosin II. This would be predicted because blebbistatin is known to act directly on myosin II.

## Discussion

The CLDN3-depletion model provides a means to investigate the specific role of the non-neural ectoderm in neural fold fusion. We demonstrated that the spinal neural folds of chick embryos button instead of zipper using live imaging analysis, consistent with a previous study (van Straaten et al., 1993). Button-like closure is also observed in small regions of the mouse midbrain (Wang et al., 2017) and hindbrain (Pyrgaki et al., 2010) as well in the spinal region of pig (van Straaten et al., 2000) and rabbit (Peeters et al., 1998b) fusing neural folds. CLDN3 depletion appears to preferentially slow the buttoning process in the spinal region but have no effect on the zippering process in the anterior cranial region or in the posterior neural pore. Our data support that CLDN3 is essential for the progression of spinal neural fold fusion by regulating non-neural ectodermal tension through pMLC activity (Fig. 7), suggesting that the buttoning process of neural fold fusion, unlike zippering, is sensitive to non-neural ectodermal tension during neural fold convergence (Fig. 7).

**Figure 7.**
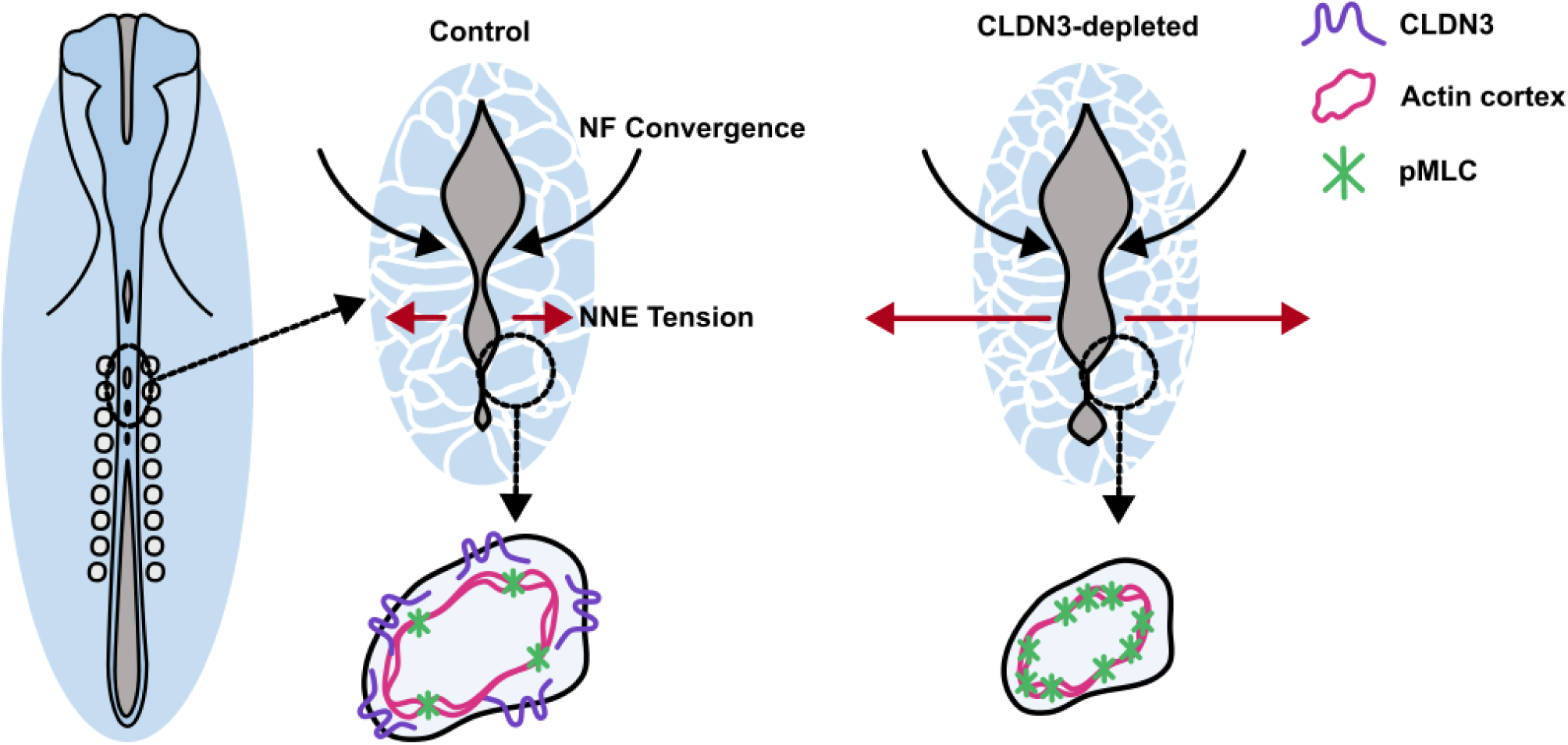
Model of the role of CLDN3 in the non-neural ectoderm during neural fold fusion. Schematic of an HH10 chick embryo undergoing neural fold fusion. Dotted circles indicate regions shown at higher magnification. Cell outlines are shown with white lines and the convergence of the neural folds with black arrows. The underlying tension in the non-neural ectoderm (NNE) is shown with red arrows. Colour legend: CLDN3: purple, Actin cortex: pink, pMLC: green, Non-neural ectoderm: blue, neural ectoderm: grey. CLDN3, through regulation of pMLC and actomyosin contraction, maintains the balance of tissue tension in the non-neural ectoderm to mediate neural fold fusion.

Although there are mechanistic differences in neural fold fusion between chick and mice, our work aligns with the NTD phenotypes observed in mice with altered expression of the transcription factors Grhl2 (Nikolopoulou et al., 2019; Ray and Niswander, 2016), and Grhl3 (Ting et al., 2003) that are expressed in the non-neural ectoderm and known to regulate expression of claudins and other cell junctional proteins (Kimura-Yoshida et al., 2015; Senga et al., 2012). Both Grhl2 and 3 regulate actomyosin dynamics (Jaffe and Niswander, 2021; Nikolopoulou et al., 2019), and Grhl2 loss leads to disorganized F-actin staining, wider and less-elongated cells, and increased lateral tissue tension. Although chick embryos do not have an actomyosin cable at the neural fold edge, CLDN3depletion decreased F-actin signal at bicellular junctions (Legere et al., 2024). While the shapes of the cells at the neural fold edges are different between mice and chick, CLDN3 depletion also leads to cell shape and size changes, with smaller apical cell areas observed on cells closest to the neural fold edge. CLDN3 depletion also led to increase tension in the non-neural ectoderm. Comparing the Grhl2-overexpression mouse model and the CLDN3-depletion chick model suggests that the functions of CLDN3 and other proteins regulated by Grhl2 in the non-neural ectoderm are conserved, although mechanisms of neural fold fusion are not. Our work contributes to a larger understanding about functions of CLDN3 in regulating the cytoskeleton to mediate tension as well as the critical role of the non-neural ectoderm in neural fold fusion.

There are several different theories to explain where force is generated during neural tube development. It was previously shows that the dorsal neural ectodermal tissue becomes stiffer during this process (Handler et al., 2023; Zhang et al., 2019), and the ability of the neural ectoderm to generate force during neural fold fusion has been observed in the dorsolateral hinge points (Roellig et al., 2022) and measured at the RCNP (Maniou et al., 2024). In the zippering chick posterior neuropore a pushing force along the anterior-posterior axis from the neural tube and notochord drives elongation and neural fold fusion (Xiong et al., 2020). We found that the non-neural ectoderm is less tense than the neural ectoderm. This feature was previously observed in *Xenopus* (Christodoulou and Skourides, 2022; Wiebe and Brodland, 2005) and led to the hypothesis that the non-neural ectoderm is passive to the forces generated by the neural ectoderm. However microsurgical experiments definitely show that the non-neural ectoderm is required for neural fold fusion (Alvarez and Schoenwolf, 1992), specifically the medial region of the non-neural ectoderm that is part of the neural folds (Hackett et al., 1997). Further evidence of the importance of the non-neural ectoderm in neural fold fusion is that depletion of proteins of Grhl transcription factors in mice (Ray and Niswander, 2016; Ting et al., 2003) and CLDN3 in chick (Legere et al., 2024) that are restricted to the non-neural ectoderm cause NTDs. Further evidence for the importance of tissue tension in neural tube closure is that preventing a loss of tension in the extra-embryonic and vitelline membrane in chick embryos causes defects in elongation and neural tube development in chick embryos (Kunz et al., 2023). Similarly in the *Xenopus* increasing non-neural ectoderm tension causes NTDs (Christodoulou and Skourides, 2022). Similar to our results with blebbistatin, decreasing tissue tension also causes NTDs in chick (Kunz et al., 2023) and *Xenopus* (Christodoulou and Skourides, 2022). Intriguingly, neural fold fusion in the spinal region of chick embryos was the most sensitive to increasing tension (Kunz et al., 2023). Although we were unable to fully evaluate whether the increased non-neural ectodermal tension following CLDN3 depletion caused NTDs in chick embryos, we were able to see a rescue in a subset of CLDN3-depleted embryos with the myosin II inhibitor blebbistatin. Blebbistatin has also been shown to partially rescue NTDs caused by RhoA inhibition, Zic2^ku/ku^ and Grhl2^Axd/Axd^ mutant mice (Escuin et al., 2023, 2015; Nikolopoulou et al., 2019). Therefore, based on our data and the work of others, we hypothesize that a precise balancing of epithelial tissue tension in the non-neural ectoderm is required to permit neural fold fusion. Balancing non-neural ectodermal tension during buttoning may be more important as there is less additional force being generated from the zippering point (Maniou et al., 2021).

Our data is not the first to suggest a relationship between tight junctions and regulation of the cytoskeleton and tissue tension. Tight junction components have been shown to interact with RhoGTPases or regulating proteins, such as ARHGAP12 (Tambrin et al., 2025), ARHGEF11 (Itoh, 2013), GEF-H1 (Aijaz et al., 2005; Haas et al., 2024), p115RhoGEF (Chumki et al., 2022), p114RhoGEF (Pinto-Dueñas et al., 2024) among others (Suarez-Artiles et al., 2022). Simultaneous depletion of CLDN3, 4, and 8 in chick embryos results in decreased RHOA, CDC42, and pMLC in the neural ectoderm (Baumholtz et al., 2017). In contrast RHOA and CDC42 are not decreased in the non-neural ectoderm in CLDN3-depleted embryos (Legere et al., 2024), and here we show that pMLC is increased in the non-neural ectoderm. These data suggest distinct functions for claudin family members in cytoskeletal regulation. A complete claudin knock-out Madin-Darby canine kidney (MDCK) cell line (quinKO: genes for CLDN1, 2, 3, 4, and 7 are deleted) was created to test the effects of loss of claudins in an epithelial cell line. Claudin quinKO MDCK cells have altered actomyosin organization, with highly developed circumferential actin bundles surrounding the cell junctions (Otani et al., 2019). Overexpression of CLDN3 in the quinKO cells reduces the circumferential staining (Fujiwara et al., 2022). Although we previously saw that in the non-neural ectoderm CLDN3-depletion decreases F-actin staining in the bicellular junctions, the cell junctional localization of PAR3 and PALS1 was increased (Legere et al., 2024), showing that CLDN3 is capable of organizing protein localization patterns in multiple different cell contexts. Claudin quinKO MDCK cells also have increased Myosin II accumulation at the cell junction and increased cortical tension during lumen expansion of MDCK cysts (Mukenhirn et al., 2024). Similar effects on tension were observed with loss of ZO1 and ZO2 in MDCK cells (Pinto-Dueñas et al., 2024). Tight junctions may play a role of regulating resistance to mechanical stress (Nguyen et al., 2024) and/or tissue fluidity (Skamrahl et al., 2021), both functions that require a tight control of biomechanical sensing and response. Despite limitations in determining the exact interactions between CLDN3 and pMLC, the increase in actomyosin contraction and tissue tension that we observed in CLDN3-depleted embryos highlights an important function for CLDN3 that should be further investigated.

## Materials and Methods

### Chick embryo collection and culture

Chick embryos were collected from white leghorn chick eggs (Ferme Gms, Saint-Roch-de-l’Achigan) incubated at 38.5 °C until the desired stage following the Hamilton and Hamburger (HH) staging guide (Hamburger and Hamilton, 1951). All embryos were handled according to the Canadian Council of Animal Care guidelines.

Chick embryos were cultured with filter papers on agar-albumin dishes following standard protocols (Chapman et al., 2001) or for some experiments included in Fig. 6 (A, B) embryos were cultured using the Cornish Pasty method (Nagai et al., 2011). For assessment of neural tube development, embryos were incubated at 38.5 °C for ∼24 hrs until gastrulation stage HH4 and cultured another ∼24 hrs until ∼HH12 when most of the neural tube development was complete. To deplete CLDN3, embryos were treated with the GST-tagged C-CPE variant C-CPE^LDR^ (LDR) (400 µg/mL) (Sonoda et al., 1999; Veshnyakova et al., 2012) that binds and removes CLDN3 from tight junctions or molarity matched GST (266 µg/mL) at HH4 and treated with Calyculin A (Sigma #208851) and Blebbistatin (Sigma #B0560) after ∼8 hrs of culture when embryos were HH7-8 (at the start of neural fold fusion). Calyculin A and Blebbistatin solutions were made with dimethyl sulfoxide (DMSO). An equal volume of DMSO used in 5µM blebbistatin and 0.025 µM Calyculin A (0.05% of final volume) was added to GST and LDR treatment. Embryos were imaged using a Zeiss Stereomicroscope Discovery V8.

### Chick embryo microsurgeries

Chick embryos were sandwiched between two filter papers and cultured on agar-albumin dishes ventral side down until reaching the required stage. The vitelline membrane was peeled back over the position of microsurgery, or over an equivalent position on non-surgically treated controls. Using an eyebrow hair as a knife, a piece of neural tube and some extra-embryonic tissue approximately the width of one somite was removed from the embryo at specific positions. To cut the embryo at the first neural fold contact point the embryos were cut at stage HH8. To cut the embryo during late neural fold fusion in the mid-spinal region the embryos were cut at ∼HH9+, and to cut the embryo at the posterior neuropore the embryos were cut at ∼HH11. Once cut the embryos were placed back into the incubator and cultured until reaching HH12, or HH14 to assess posterior neuropore closure.

### Chick embryo live imaging

Chick embryos were incubated at 38.5 °C for ∼24 hrs and collected on filter papers at HH4 and cultured on agar-albumin dishes containing either LDR (400 µg/mL) or molarity matched GST (266 µg/mL) until the start of NF fusion (∼HH8). At ∼HH8 embryos were moved to a glass bottom imaging dish (MatTek: P35G-1.5-10-C) coated in a thin (∼ 500 µL) agar-albumin layer with on either LDR (400 µg/mL) or molarity matched GST (266 µg/mL). Simple saline was dropped on the corners of the filter paper and the embryo was placed in a humidified platform and live imaged with a Zeiss Axio Observer A1 at 38.5 °C. Embryos were imaged with an AxioCam MR R3 camera for up to 8 hrs with transmitted light halogen lamp (∼2.5 V) and minimal exposure time (100-200 msec) with 5- or 15-minute intervals and Z-stacks set up to center on NF edges with ∼ 5 µm steps. Embryos were either imaged with Fluor 10X/0.5 M27 or Plan-Apochromat 20X/0.8 objectives. Maximum intensity projections (MIPs) were made of the videos and images were automatically adjusted for brightness and contrast with ImageJ. Landmarks at the center of the RCNP were used for measuring the length between the fusing NFs and bounding rectangles were used to compare only widths between embryos. To measure the rate of zippering in the midbrain (MB) region of the embryo, the distance between the zippering point and the left somite was measured.

### Chick embryo immunofluorescence

Chick embryos were incubated at 38.5 °C for ∼24 hrs until collected on filter papers at HH4 and cultured on agar-albumin dishes on either LDR (400 µg/mL) or molarity matched GST (266 µg/mL) for 12 hrs (∼HH9). In experiments testing the effect of 5µM blebbistatin or 0.025 µM Calyculin A added at stage ∼HH8, embryos were cultured for 3 hrs after treatment before fixing. Embryos were collected and fixed in the appropriate fixative for 1 hr RT (4% PFA: ZO1, F-actin (Phalloidin)) (10% TCA: pMLC, ZO1). For pMLC immunofluorescence experiments embryos underwent antigen retrieval with 10mM Sodium Citrate for 25 mins at 100 °C. After PBS washes, embryos were blocked in 10% Normal Goat Serum (NGS) with 0.3% Triton X-100 in PBS. Embryos were incubated with primary antibodies (ZO1: Invitrogen #33-9100, 1:200) (pMLC: Cell Signalling #3674S, 1:50) O/N at 4 °C. After PBS washes embryos were incubated with secondary antibodies for 1 hr RT, and if required Phalloidin (Invitrogen #A22287, 1:250) and/or DAPI (Fisher #D1306, 1:1000) was added in this step, then washed with PBS and placed on microscope slides with FluorSave (Millipore #34578920ML). Embryos were imaged with an LSM 880 Elyra AxioObserver with either a Plan-Apochromat 20X/0.8 M27 or Plan-Apochromat 63X/1.4 Oil DIC M27 objectives.

### Analysis of immunofluorescence images

Images were analyzed using ImageJ (Schindelin et al., 2012) and surface projections were made with the macro SurfacePeeler (github.com/DaleMoulding/SurfacePeeler). Mean grey intensity was measured and the ratio of pMLC to ZO-1 mean grey value was calculated. One embryo was removed from the analysis after outlier analysis. ZO1 signal was used to define cell membrane regions and mean grey intensity was measured within the membrane region and the inverse (cytoplasm). For graphical representations the average pMLC/ZO-1 ratio per experiment was used as a normalization factor, however statistical analysis was performed on raw data.

Surface projections were segmented on using the epyseg algorithm of Tissue Analyzer (Aigouy et al., 2016) (github.com/baigouy/tissue_analyzer) using ZO-1 signal. Segmentation was hand-corrected, and the masks were used in ImageJ to measure cell properties. The images were rotated so that the anterior-posterior axis was parallel to the edge of the image. The edges of the neural folds (NFs) were traced and the distance of cells from the edge of the NF was calculated using the coordinates of the NF lines and the coordinates of the cell centroid. The distance between the NFs was also calculated for every cell. Cells with areas above or below the top/bottom 5% were removed from the analysis. For assessment of cell measurements by distance of the cell from the NF the data was binned in increments of 10µms. For assessment of cell measurements by distance between the NFs, the data was binned into 0µm, 0 <= 10 µm, and > 10 µm. Analysis, statistical tests, and graphs were done in R.

### Laser ablations

Laser ablations were performed through an on-site collaboration with Dr. Gabriel Galea’s laboratory at University College London. Deklab chicken eggs were incubated at 38.5 °C for ∼24 hrs and collected on filter papers at stage HH4. Embryos were cultured at 38.5 °C on either LDR (400 µg/mL) or molarity matched GST (266 µg/mL) until the start of NF fusion (∼HH8). Embryos were removed from the incubator immediately prior to imaging, flipped dorsal side up and the vitelline membrane over the area to be ablated/imaged was removed using a sharpened tungsten needle. Embryos were rinsed with simple saline then ∼20ul of CellMask 647 (Invitrogen #C10046, 1:100) was added dropwise to the neural tube and incubated for 15 min in 38.5 °C. CellMask was washed 3X with PBS and placed into the 38.5 °C incubator of LSM880 Examiner. Embryos were imaged using a 20X NA:0.7 C Epiplan-Aprochromat objective (1024X1024, 9.63 pixels/µm). 500-pixel long lines (∼50 µm) were ablated with a SpectraPhysics Mai Tai eHP DeepSee multiphoton laser (710 nm λ, 100% laser power). Images were captured immediately prior and after ablation (0.96 sec per image).

### Analysis of laser ablations

Final analysis was restricted to embryos up to stage HH10 and on images without air bubbles (GST: 10 embryos, 53 images, LDR: 8 embryos, 62 images). Images were analyzed using ImageJ (Schindelin et al., 2012) by selecting out the region around the ablation, running the plugin pureDenoise with default parameters (Luisier et al., 2010), running StackReg with rigid body correction (EPFL) and selecting out the time points immediately before and after ablation. Landmarks that appear in both pre- and post-ablation on either side of the center part of the ablation were selected and distances were measured three times and averaged. A bounding rectangle of the distance was used to measure the height of recoil and the difference between heights of pre- and post-ablation was measured to get the final recoil. Measurements were averaged across embryos based and Student’s T-tests or paired Student’s T-tests when applicable were used to assess significance

## Supplemental material

**Supplemental Figure 1.**
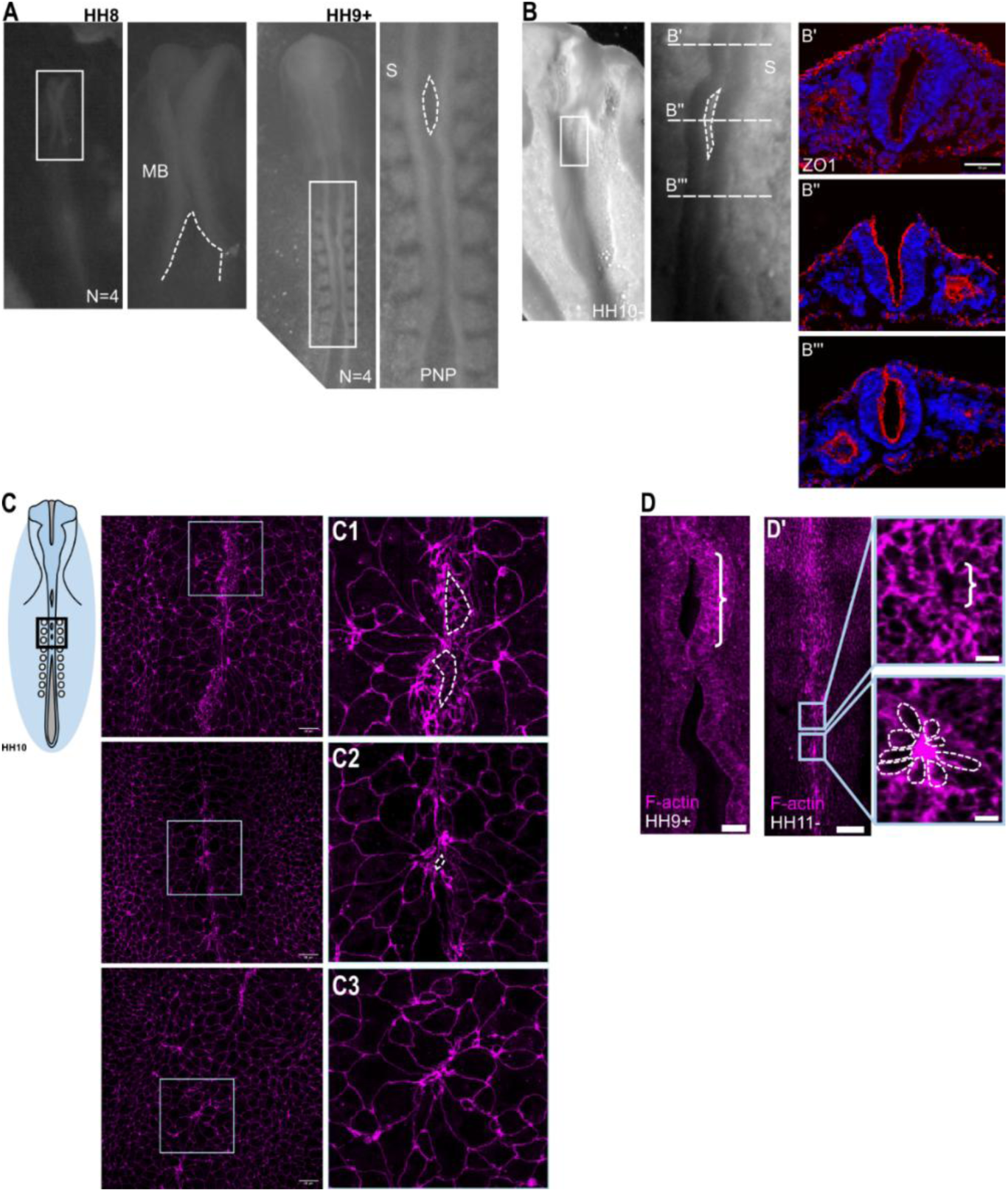
Control embryos during neural fold fusion. A) Representative images of *in ovo* embryos at HH8 (n=4) and HH9-HH10 (n=4). Neural folds (NFs) are outlined with white dotted lines. White boxes show regions shown in adjacent higher magnification image. B) HH10 embryo cryosectioned prior to immunofluorescence (red = ZO1, blue = DAPI) (n=3). White box shows region shown in adjacent higher magnification image. White dotted lines outline the RCNP. White dashed lines show approximate region show in transverse sections in B’,B’’,B’’’. Sections are ∼200µm apart. C) Maximum Intensity Projections (MIPs) of representative images from HH10 (n=3) embryos imaged after immunofluorescence with ZO-1 antibody (magenta). White boxes show regions in zoomed images and white dotted lines outline gaps between NFs of C1) open gaps, C2) fusing gaps, and C3) fused gap. D) Representative images of HH8 (n=3) and D’) HH10 (n=4) embryos stained with phalloidin (F-actin = magenta). Brackets show regions between open NFs. White dotted lines outline cells surrounding fusion. Labels: MB = midbrain, S = somite, PNP = posterior neuropore. Scale bars: B,D)5 µm, C) 20µm

**Supplemental Figure 2.**
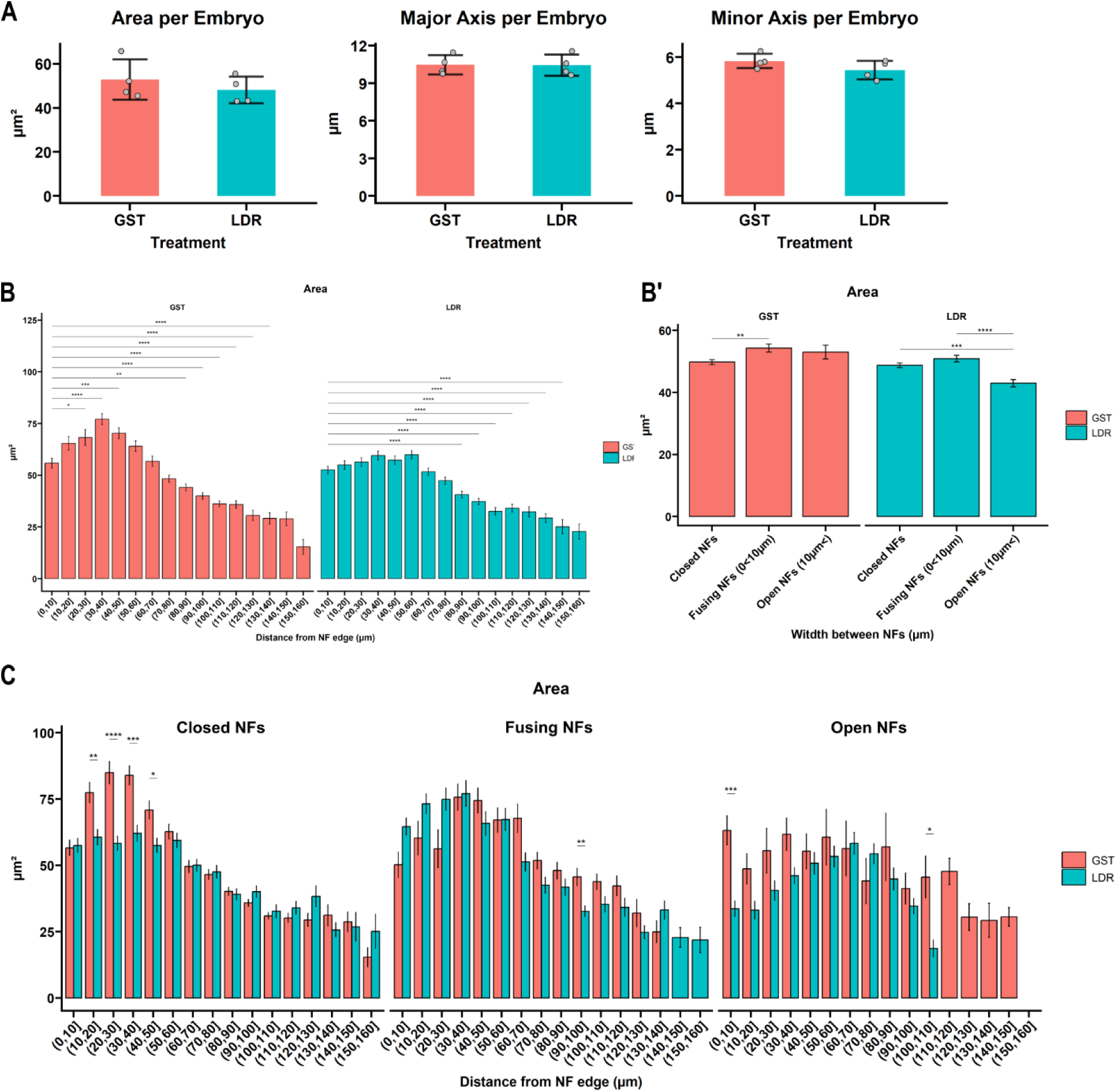
Additional cell measurements in the non-neural ectoderm during neural fold fusion. A) Quantification of the apical cell area (µm^2^), major axis (µm), and minor axis (µm) averaged per embryo comparing control (GST; pink) and CLDN3 depletion (LDR; blue). Bars show the mean, error bars plot the standard error, and grey dots show the average measurement for each embryo. One-way ANOVA of apical cell area (µm^2^) by B) cell distance bins per treatment and B’) neural fold (NF) width bins per treatment. Significance was determined with Tukey’s post-hoc analysis and comparisons on B) are shown between the 0µm – 10µm bin and every other distance. C) Analysis of cells binned by distance from the NF edge and width between NFs (Closed NFs = 0 µm, Fusing NFs = 0 µm <= 10 µm, Open NFs = > 10 µm). Statistical significance between control (GST; pink) and CLDN3-depleted (LDR; blue) in each bin was assessed with Student’s T-test with multiple comparisons with Benjamini-Hochberg (False Discovery Rate) p-value correction. **** p < 0.001, *** p < 0.005, ** p < 0.01, * p< 0.5

**Supplemental Figure 3.**
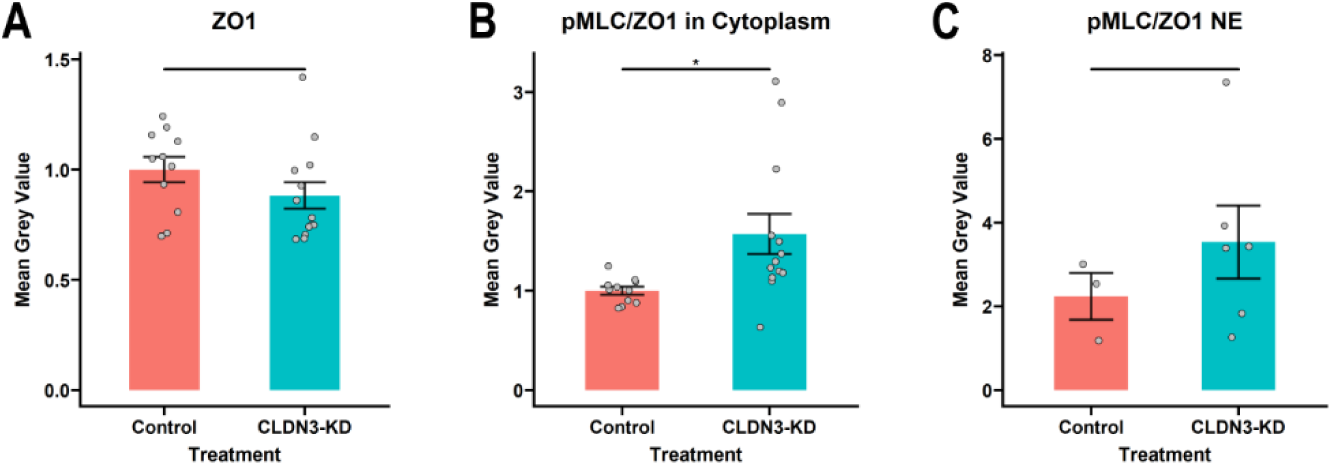
Further analysis of pMLC expression in chick. A) Quantification of ZO1 fluorescence (GST: n=12; LDR: n=11), B) ratio of pMLC/ZO1 in the cytoplasmic region of cells (GST: n=12; LDR: n=11), and C) ratio of pMLC/ZO1 in the neural ectoderm (GST: n=3; LDR: n=6) between control (pink) and CLDN3-depleted (CLDN3-KD) (blue). Bars show the mean, error bars plot the standard error, and grey dots show the average measurement for each embryo. Statistical significance was evaluated with Student’s T-test. * p< 0.5.

**Supplemental Figure 4.**
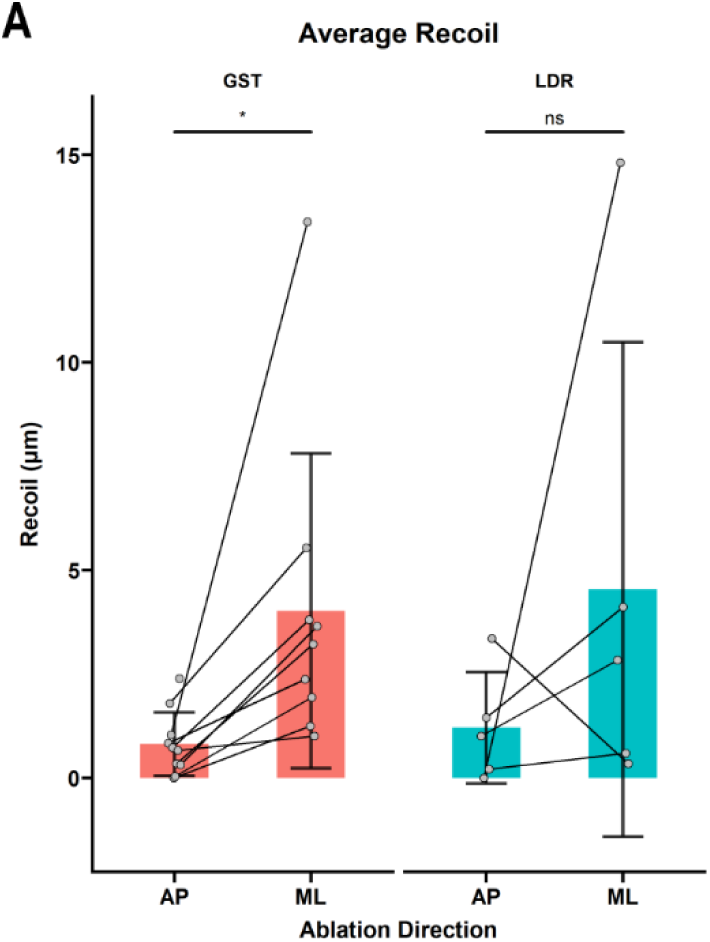
Tension in the neural ectoderm is anisotropic. A) Quantification of recoil in the neural ectoderm (NE) of control (WT) (n=9) and CLDN3-depleted (LDR) (n=5) embryos. Ablations in the medial lateral (ML) and anterior posterior (AP) direction were compared. Points represent the average per embryo, and lines connect measurements made on the same embryo. Statistical significance was assessed with paired Student’s T-test between ML and AP directed ablations. * p< 0.5.

**Supplemental Figure 5.**
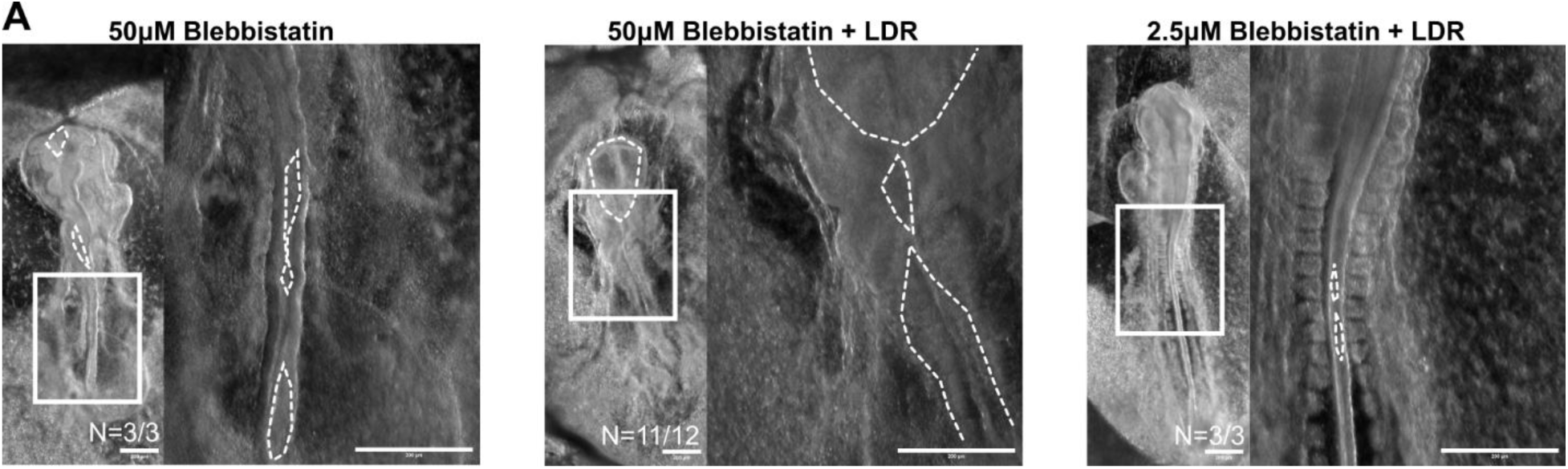
Embryos treated with high and low dose blebbistatin. A) Representative images of 50µM Blebbistatin (n=3), 50µM Blebbistatin + 400µg/mL LDR (n=12), and 2.5µM Blebbistatin + 400 µg/mL LDR (n=3) after 24 hr of culture, treated with blebbistatin at HH8. White boxes illustrate region of image shown in zoomed image; white dotted lines outline open neural tube defects (NTDs). Scale bars: 200µm

## Abbreviations

ANP: anterior neuropore
AP: anterior - posterior
C-CPE: C-terminal of *Clostridium perfringens* enterotoxin
CLDN3: Claudin-3
DLHP: dorsolateral hinge point
GST: glutathione S-transferase
LDR: variant of C-terminal of *Clostridium perfringens* enterotoxin
MB: midbrain
ML: medial – lateral
NE: neural ectoderm
NNE: non-neural ectoderm
NF: neural fold
NTD: neural tube defect
pMLC: phosphorylated myosin light chain
PNP: posterior neuropore
RCNP: rhombocervical neuropore
ZO1: zona occludens 1
ZP: zippering point

## Data availability statement

Data is available upon request

## Acknowledgments

We thank Dr. I. Gupta, Dr. R. Rozen, Dr. D. Reinhardt, and Dr. D. Dufort for their feedback and discussions throughout the project, as well as members of the Ryan and Gupta labs, specifically to V. Murugapoopathy for review of the manuscript, and Z. Budhwani for excellent eyebrows used for microsurgery. This work was supported by a Discovery Grant from the Natural Science and Engineering Research Council of Canada (NSERC) (RGPIN-2023-04809) and funding from Fondation des étoiles (AKR). EAL is a recipient of a RI-MUHC studentship award, a Faculty of Medicine and Health Sciences Arthur Judson studentship, a masters Alexander Graham Bell and doctoral studentship (546885) from NSERC, and a doctoral studentship from Fonds de recherche du Québec – Santé (FRQS) (292904). This work was supported by the Natural Sciences and Engineering Research Council of Canada (AKR). AKR is a member of the RI-MUHC, which is supported in part by the FRQS. Thank you to Dr. G. Galea, Dr. N. Greene, Dr. A. Copp and their labs for hosting me for laser ablation experiments, with a special thank you to The Company of Biologist for funding, and Dr. G. Galea for his effort, time, and helpful discussions. GLG acknowledges support from the Wellcome Trust (211112/Z/18/Z), Royal Society (RG\R2\232082) and Leverhulme Trust (RPG-2024-147). This work was performed with the support of the RI-MUHC Molecular Imaging Platform.

